# Conservation implications of elucidating the Korean wolf taxonomic ambiguity through whole-genome sequencing

**DOI:** 10.1101/2023.02.24.529912

**Authors:** Germán Hernández-Alonso, Jazmín Ramos-Madrigal, Xin Sun, Camilla Hjorth Scharff-Olsen, Mikkel-Holger S. Sinding, Nuno F. Martins, Marta Maria Ciucani, Sarah S. T. Mak, Liam Thomas Lanigan, Cecilie G. Clausen, Jong Bhak, Sungwon Jeon, Changjae Kim, Kyung Yeon Eo, Seong-Ho Cho, Bazartseren Boldgiv, Gankhuyag Gantulga, Zunduibaatar Unudbayasgalan, Pavel A. Kosintsev, Hans K. Stenøien, M. Thomas P. Gilbert, Shyam Gopalakrishnan

## Abstract

The taxonomic status of the now likely extirpated Korean Peninsula wolf has been extensively debated, with some arguing it represents an independent wolf lineage, *Canis coreanus*. To investigate the Korean wolf genetic affiliations and their taxonomic implications, we sequenced and analysed the genomes of a historical Korean wolf dated to the beginning of the 20th century, and a captive wolf originally located at the Pyongyang Central Zoo. Our results indicated that the Korean wolf bears similar genetic ancestry to other regional East Asian populations, therefore suggesting it is not a distinct taxonomic lineage. We identified regional patterns of wolf population structure and admixture in East Asia with potential conservation consequences in the Korean Peninsula and on a regional scale. We find the Korean wolf has similar diversity and inbreeding to other East Asian wolves. Finally, we show that, in contrast to the historical sample, the captive wolf is more genetically similar to wolves from the Tibetan Plateau, hence, Korean wolf conservation programs might not benefit from the inclusion of this specimen.

## Introduction

Species and subspecies ambiguity have been a common problem in canid taxonomy due to their wide and continuous distribution, their long-distance mobility across topographic barriers, the extended gene flow among populations, and even, the hybridization among closely related species (Gopalakrishnan et al. 2018; Nowak 1995; Nowak 2003; Pilot et al. 2006; Shrotriya et al. 2012; Wayne et al. 2003). A specific case of this ambiguity problem that has not been properly resolved is represented by the likely extirpated wolf populations from the Korean Peninsula.

The Korean wolf geographic distribution spanned all the Korean Peninsula until the middle of the last century. However, its population was severely impacted by the Japanese incursion into Korea (1910-1945) and their industrialization program. The remaining populations were finally extirpated from South Korea during the late 20th century. The last individuals were recorded during the 90s around Paektu Mountain, located at the border between North Korea and China (Jo et al. 2018). At present, wolves are considered extirpated from the entire Peninsula according to the Red Data Book of South Korea (National Institute of Biological Resources, 2012). However, there is very limited information about the conservation status of wolf populations in North Korea.

Korean wolf populations have been subject to several taxonomic changes since the late 19th century. At first, Korean wolves were included in the original definition of *Canis chanco* proposed by Gray (1863) together with populations from the Chinese Tartary (Central China, Northern China, and Mongolia). Later, Mivart (1890) suggested that *C. chanco* and the wolves from Tibet, *Canis laniger* Hogdson, 1847, were probably the same taxonomic group, but also, that both forms were varieties of the common grey wolf, *Canis lupus* Linnaeus, 1758, changing them to a subspecies category. The status of the Korean wolf populations was not revised until 1923, when the Japanese zoologist Yoshio Abe suggested that the Korean wolves were morphologically different enough from the continental populations to be considered a different lineage: *Canis coreanus* (Abe 1923). The British zoologist Reginald Pocock rejected such proposal, including the Korean wolves in the form *Canis lupus laniger* (synonym of *C. l. chanco*) together with populations from the “Chinese Tartary, Thian Shan, Kashmir, Tibet, Mongolia, and China” (Abe 1936).

In 2005, Wozencraft reasserted the subspecies condition of *C. l. chanco*, including it as one of his synonyms *C. coreanus* Abe, 1923. Importantly, the *C. l. chanco* subspecies by itself is a well-known case of taxonomic ambiguity that has been largely discussed (*cf*. Aggarwal et al. 2003; Aggarwal et al. 2007; Joshi et al. 2020; Shrotriya et al. 2012). By that time, the *C. l. chanco* subspecies distribution was considered to cover a large extension of central and east Asia, as suggested by Sokolov and Rossolimo (1985). Later, genomic data confirmed the high degree of divergence between Himalayan and Tibetan Plateau wolf populations (Aggarwal et al. 2003), as well as their unique adaptations to high altitude environments (Werhahn et al. 2018). Then, in 2019, the IUCN/SSC Canid Specialist Group recommended the use of the name *Canis lupus chanco* for those populations restricted to the Himalaya range and the Tibetan Plateau (Alvares et al. 2019). As a consequence, the rest of Asian wolf populations included before in *C. l. chanco* were classified as *C. l. lupus*. Although these taxonomic changes included the Korean wolf population, its validity has not been tested using genomic data.

The first time that a Korean wolf was included in a molecular analysis was by Ishiguro et al. (2009; 2010), where the mitochondrial control region (751bp) was used to study the extinct Japanese wolf (*C. l. hodophilax*) and Ezo wolf from Hokkaido (*C. l. hattai*). The Korean wolf sample used in that study is described as a *C. l. chanco* and it was collected from the Hasebe Collection at the University of Tokyo Museum. In the results presented by Ishiguro et al. (2009; 2010) the Korean wolf seems to be closely related to Chinese and Mongolian wolves, as would be expected based on geography. However the variation in the mitochondrial control region has limited resolution, hence additional work including autosomal variation is needed to truly evaluate the affiliation of original Korean wolves.

Here, we generated whole-genome sequencing data from a captive wolf, putatively North Korean, as well as a historical wolf from South Korea, to an average depth coverage of 25.29× and 7.25× respectively (Table S1). Additionally, we resequenced 10 wolves from poorly characterised populations in Russia, Mongolia and Kazakhstan, and a Mongolian deceased captive wolf from Zoo Zürich to an average depth coverage of 5-7×, to obtain a better representation of Asian wolf populations (Table S1). The historical Korean wolf genome was obtained from a female specimen housed at the Kyungpook National University Natural History Museum, Gunwi-gun, South Korea. This individual dates to the beginning of the 20th century (before 1945), and given its provenance, it likely represents one of the last wolf populations in South Korea. The modern genome was obtained from a female captive wolf, originally located in the Pyongyang Central Zoo, North Korea, and subsequently transferred to the Seoul Grand Park in 2005. Her wild origin is ultimately uncertain. Using these samples, we aimed at resolving the taxonomic ambiguity of the Korean wolf, searching for the closest related continental wolf populations and analysing their level of genetic differentiation. Moreover, we endeavor to track the potential origin of the captive wolf to confirm if its lineage came from inside or outside of the Korean Peninsula.

## Materials and Methods

### Description of wolf samples

Three sup-samples of the historical Korean wolf (HKW) were collected from the skin of a female stuffed specimen housed at the Kyungpook National University Natural History Museum, Gunwi-gun, South Korea. Blood samples were taken from a female captive wolf (PZW) at the Seoul Grand Park that was transferred from the Pyongyang Central Zoo, North Korea on April 14, 2005, through animal exchange between South and North Korea. Finally, a tissue sample was obtained from a deceased captive Mongolian wolf (A2) (*Canis lupus chanco*, Local ID: ZURICH / A70094) Zoo Zürich.

Muscle samples from the modern specimens 750021A and 750115A were provided by the Yekaterinburg Museum, Russia. A further skin sample (MW486) and extracts (MW524, MW536, and MW538) were also collected from Russian wolf populations and provided by collaborators. Samples from muscle, cheek tissue, and skin corresponding to MW561, MW574, and MW588 were obtained from contemporary Mongolian wolves.

These data were generated as part of a broader study funded by the Norwegian Environment Agency that aims at looking at the relationship of modern and historic Norwegian wolves to other Eurasian wolves (Stenøien et al.)

### Laboratory processing of modern samples

DNA from modern wolf PZW (SGP-824) was extracted using the DNeasy Blood & Tissue kit (Qiagen) according to the manufacturer’s recommendations. DNA quality was assessed by running 1 μL on the Bioanalyzer system (Agilent) to ensure size and analysis of DNA fragments. The concentration of DNA was assessed using the dsDNA BR assay on a Qubit fluorometer (Thermo Fisher Scientific). DNA was converted into double stranded blunt-end libraries with BGI-specific adapters (Mak et al. 2017) using the BEST protocol (Carøe et al. 2018). Libraries were sequenced on a BGISeq 500 plataform using 100 base pair paired-end read chemistry.

DNA from the modern Mongolian wolf (A2) was extracted using a DNeasy Blood & Tissue Kit (Qiagen) following the manufacturer’s protocol. DNA was converted into double stranded blunt-end libraries with Illumina-specific adapters REF using the NEBNext DNA Sample Prep Master Mix Set 2 (E6070S - New England Biolabs Inc., Beverly, MA, USA) following the manufacturer’s protocol. Libraries were sequenced on a Illumina HiSeq 2500 platform using 100 base pair paired-end read chemistry.

DNA from modern wolves samples (MW486, MW524, MW536, MW538, MW561, MW574, MW588, 750021A, 750115A) was extracted and prepared for sequencing in the DNA laboratories at the Globe Institute, University of Copenhagen, using a Kingfisher duo prime extraction robot. DNA extracts were sent to BGI Denmark for library building and sequencing. The samples were sequenced in a DNBseq platform in PE150 mode.

### Laboratory processing of historical samples

The HKW sample was processed under strict clean laboratory conditions at the Globe Institute, University of Copenhagen. Three tissue samples were placed into 3 eppendorf tubes – 1, 2 and 3 – and washed with diluted bleach, ethanol and ddH2O, following Boessenkool et al. (2017). The material was processed following Gilbert et al. (2007) DNA extraction protocol. Additional treatment with phenol chloroform was performed following Carøe et al. (2018). The supernatant was then purified using a modified PB buffer and eluted using 2 washes in 18 μl buffer EB (QIAGEN) - with 3 min of incubation time at 37°C (Dabney et al. 2013). The concentration of each extract was checked on a Qubit (ng/μl). BGI libraries were built using 10-20 μl of extract in a final reaction volume of 50 μl following the Santa Cruz Single Stranded protocol (Kapp et al. 2021), and using the “single-tube” library building protocol BEST (Carøe et al. 2018). Library index amplifications were performed using PfuTurbo Cx HotStart DNA Polymerase (Agilent Technologies) in 50 μL PCR reactions that contained 5 μL of purified library, 0.1 μM of each forward (BGI 2.0) and custom made reverse primers (Mak et al. 2017). The PCR cycling conditions were: initial denaturation at 95°C for 2 mins followed by 20 cycles of 95°C for 30 s, 60°C for 30 s, and 72°C for 2 mins, and a final elongation step at 72°C for 10 mins. Amplified libraries were then purified using 1.8x ratio of MagBio beads to remove adaptor dimers and eluted in 30 μL of EBT (QIAGEN) after an incubation for 5 min at 37°C. Amplified libraries were sequenced at Clinomics. Inc with a SE100 mode, and at BGI China with a SR100 mode.

### Dataset

The dataset used for this study includes 74 canid whole-genomes: one Andean fox (*Lycalopex culpaeus*) (Auton et al. 2013), two coyotes (*Canis latrans*) (Gopalakrishnan et al. 2018; vonHoldt et al. 2022), 51 wolves (*Canis lupus*) (Hennelly et al. 2021; Ramos-Madrigal et al. 2021; Niemann et al. 2021; Sinding et al. 2020; Sinding et al. 2018; Fan et al. 2016; vonHoldt et al. 2016; Wang et al. 2016; Wang et al. 2013; Freedman et al. 2014; Zhang et al. 2014), and 20 dogs. (Sinding et al. 2020; Kolicheski et al. 2017; Marchant et al. 2017; Metzger et al. 2017; Marsden et al. 2016; Wang et al. 2016; Decker et al. 2015; Freedman et al. 2014; Auton et al. 2013; Kim et al. 2012; Lindblad-Toh et al. 2005) Among the wolf samples, 40 are reference genomes mainly representing Asian populations; five correspond to Pleistocene wolves from Siberia, nine correspond to wolves resequenced for this study representing wolf populations in Asia, and two are the Korean wolves: a modern captive wolf (PZW) from the Seoul Grand Park in South Korea, and a 20th century stuffed Korean wolf (HKW) from the Kyungpook University Museum. Finally, all dog genomes included here are already published and they were selected to mainly represent Asian breeds (Table S2).

### Data processing

Sequence reads obtained from BGI were mapped to the wolf reference genome (Gopalakrishnan et al. 2017) using PALEOMIX v.1.2.13.3 BAM pipeline (Schubert et al. 2014). The wolf reference genome was generated from a highly inbred Swedish wolf, which mitigates issues with reference mapping bias. In brief, adapter trimming was performed using AdapterRemoval v.2.2.0 (Schubert et al. 2016) and only reads with a minimum length of 25bp were kept. Trimmed reads were mapped to the reference genome using BWA v.0.7.16a backtrack algorithm (Li and Durbin 2009) disabling the use of a seed parameter. PCR duplicates were identified and removed using Picard MarkDuplicates (Broad Institute 2016) and, finally, local realignment around indels was performed using GATK v.3.8 3 IndelRealigner module (McKenna et al. 2010).

To evaluate the substitution patterns in the sequences of the HKW DNA we used mapDamage v.2.0.9 (Jónsson et al. 2013). The aligned sequences resulting from the mapping process were used as input with the default parameters. This step allowed us to authenticate the historical condition of the analysed sample.

### Multidimensional scaling plot

We performed SNP calling by randomly sampling a read for every site for all the samples in our dataset using the option -doHaploCall from ANGSD v.0.931 (Korneliussen et al. 2014). This option allows sampling a random read from each site and each sample instead of performing genotype calling, which allows the incorporation of low to medium depth of coverage samples. We used the following parameters: doCounts 1 -minMinor 2 -maxMis 7 –C 50 -baq 1 -uniqueOnly 1 -remove_bads 1 -only_proper_pairs 1 -skipTriallelic 1 - doMajorMinor 1. Additionally, bases with quality lower than 20 and mapping quality lower than 30 were discarded. Transitions were removed to avoid aDNA damage that could be found in historical samples. A MAF filter of 0.01 was applied given a final SNPs dataset of 3,284,758 transversion sites. Finally, we restricted the analysis to the scaffolds with at least 1 Mb in size.

Using the previously mentioned SNPs dataset, we estimated pairwise distances between the samples, and generated a multidimensional scaling (MDS) plot using Plink v.1.90 (Chang et al. 2015). We estimated an MDS plot including all samples except for the outgroups, and a second one restricting to wolves.

### Admixture analysis

In order to estimate the ancestry components in the historical Korean wolf (HKW) and the wolf (PZW) from South Korean Zoo included in our dataset, we used the previously described SNPs panel and ADMIXTURE v.1.3.0 (Alexander et al. 2009). Outgroups were excluded from this analysis. We ran ADMIXTURE assuming 2 to 10 (K2-K10) ancestry components. Ten replicates for each K value were performed, and the replicate with the best likelihood value was selected. R library Pophelper v.2.3.1 (Francis 2017) was used to visualize the admixture plots.

### Neighbor-joining tree

A neighbor-joining tree (NJ) was estimated based on a distance matrix constructed in Plink

1.90. The same SNPs panel described before was used as input to construct the distance matrix. The tree was estimated using R library *ape* (Paradis and Schliep 2019) and Interactive Tree Of Life (iTOL) v4 online tool (Letunic and Bork 2019) was used to visualize it.

### Maximum likelihood nuclear genome phylogeny

A maximum-likelihood nuclear genome phylogeny was inferred to explore the evolutionary relationships among the samples from our dataset. The Andean fox and two coyotes were included as outgroups. For each genome, ANGSD v.0.931 was used to generate genomic consensus sequences using the wolf reference genome (“-dofasta2” option). Then, 1000 independent phylogenetic trees were estimated with RAxML-ng v.0.9.0 (Kozlov et al. 2019) under the GTR+G evolutionary model, using 1000 random regions of 5000 bp. All gene trees were concatenated to generate a species tree using ASTRAL-III (Zhang et al. 2018) which was visualized using the Interactive Tree Of Life (iTOL) v4 online tool (Letunic and Bork 2019).

### TreeMix

To explore admixture patterns in our historical Korean wolf, the software TreeMix v.1.13 (Pickrell and Pritchard 2012) was used to estimate an admixture graph. We included a subset of samples conformed by the historical Korean wolf (HKW), the Pyongyang Zoo wolf (PZW), Asian wolves that appeared close to the HKW based on the *f3*-statistics and the estimated NJ-tree, as well as wolves representing different clades on the tree (Portuguese, Xinjiang-CAN30, Indian-BH6 and Tibet-CAN9A), and Asian dog breeds, excluding other dog breeds to avoid the estimation of migration events among the different dog lineages. The final subset consisted of 16 wolves, 12 dogs and 2 coyotes used as outgroups. TreeMix was run using the previously created SNPs panel, restricting the analysis to sites without missing data (1,904,538 sites), estimating 0 to 5 migration events (-m) and grouping SNPs in windows of 500 (-K). Ten replicates were performed for each migration event and the one with the best likelihood was chosen. The final results were plotted on R using TreeMix script plotting_funcs.R.

### Outgroup f3-statistics

We calculated outgroup *f3-statistics* using the qp3pop tool from ADMIXTOOLS v.5.1 package (Patterson et al. 2012) to measure the amount of share drift between two populations since its common ancestor. For this analysis we used the coyotes as the outgroup. We estimated the shared drift between all our samples and the two Korean wolves to find its most closely related population. Higher values indicate a closer relationship due to shared drift between the two populations in the test.

### D-statistics

We used our SNPs panel to run *D*-statistics, as implemented qpDstat from ADMIXTOOLS v.5.1 (Patterson et al. 2012) to evaluate the possibility of admixture between the historical Korean wolf and dog breeds, or the captive wolf from the Seoul Grand Park (Pyongyang Zoo wolf) and dog breeds. For a given test in the form D(Outgroup, A; B, C), if the obtained *D*-statistic significantly deviates from 0 it suggests possible gene flow between A and B (D < 0) or A and C (D > 0). In all the analyses the Andean fox was used as Outgroup. Deviation from 0 was considered statistically significant when Z-score was below -3 or above 3. The significance of the test was assessed using a weighted block jackknife procedure over 1 Mb blocks.

We evaluate the following tests for both the historical Korean wolf and the captive Pyongyang Zoo wolf:

- D(Outgroup, Korean wolf; Tibetan Mastiff, Dogs)
- D(Outgroup, Pyongyang Zoo wolf; Tibetan Mastiff, Dogs)
- D(Outgroup, Eurasian wolves; Tibetan Mastiff, Shar Pei)
- D(Outgroup, Dogs; Portuguese wolf, Korean wolf)
- D(Outgroup, Dogs; Portuguese wolf, Pyongyang Zoo wolf)

### Genotype imputation

In order to have a more robust dataset for missingness-sensitive analysis, we imputed the genotype with Beagle v5.4 (Browning et al. 2018). First, site allele frequency likelihoods were generated from elven high coverage samples (>7X) using ANGSD (-doSaf 1) based on GATK’s model (-GL2), excluding transitions (-noTrans 1) and reads with mapping quality lower than 20 (-minQ 20). The wolf reference genome was used as both the ancestral (-anc) and reference (-ref) genome. The obtained genotype likelihoods were genotyped for all samples (N=71) using bcftools v1.12 (Danecek and McCarthy 2017). Scaffolds with a length less than 10kb were discarded. Beagle was then used for imputation with default parameters using a sliding window size of 40.0 cM and an overlap of 2 cM between adjacent windows. After applying a MAF filter of 0.05, we obtained a final dataset of 4,036,734 transversion sites.

### Heterozygosity per windows and inbreeding coefficient

Imputed genotypes were used to calculate heterozygosity per window using Plink v.2.3.8. Window’s size of 1Mb were defined on the first 704 scaffolds longer than 1Mb. The Plink function *--sample-counts* was implemented to estimate the number of heterozygous sites for each window, and finally, heterozygosity per window ratio was calculated (heterozygous sites/total sites) and the results were visualized using R. To estimate the inbreeding coefficient we implemented the Plink function *--het* on the same imputed genotype dataset.

## Results

We obtained whole-genome sequences for a total of 12 wolf specimens, including the captive wolf originally held at the Pyongyang Zoo (PZW), the historical Korean wolf (HKW), 5 wolves from Russia, 4 from Mongolia (one captive in a Swiss zoo), and 1 from Kazakhstan. These genomes were combined with available reference genomes, comprising whole-genome data from 51 wolves, 20 dogs, 2 coyotes and 1 Andean fox (Table S2). Sequencing data for the new and reference samples was mapped to the wolf reference genome (Gopalakrishnan et al. 2017). To account for the low coverage samples, we performed SNP calling using ANGSD by randomly sampling a read for every site in the genome. We selected variable sites with a MAF of 1% and restricted to transversion sites to avoid biases related to historical DNA damage. Our final dataset consisted of 3,284,758 transversion sites. For analyses requiring diploid genotypes, we generated a second dataset with a subset of the high coverage samples and performed genotype calling and imputation. The imputed genotypes dataset consisted of 4,036,734 transversion sites.

### Korean wolves’ population affinities

To explore the population structure in our dataset and assess how the PZW and the HKW relate to wolf and dog populations worldwide we used a multidimensional scaling (MDS) analysis. The results clearly demonstrate that the samples cluster in three main groups: dogs, highland wolves (represented by the wolves from the Tibetan Plateau), and the rest of the wolves in our dataset (Figure 1A). Dimension 1 separates dogs and wolves, while dimension 2 separates the highland wolves from the rest of the wolf populations. It is notable that the two Korean samples are not placed together, but rather the HKW clusters in the main wolf cluster near to the east Asian wolves, while the PZW clusters together with highland wolves from the Tibetan Plateau.

**Figure 1.**
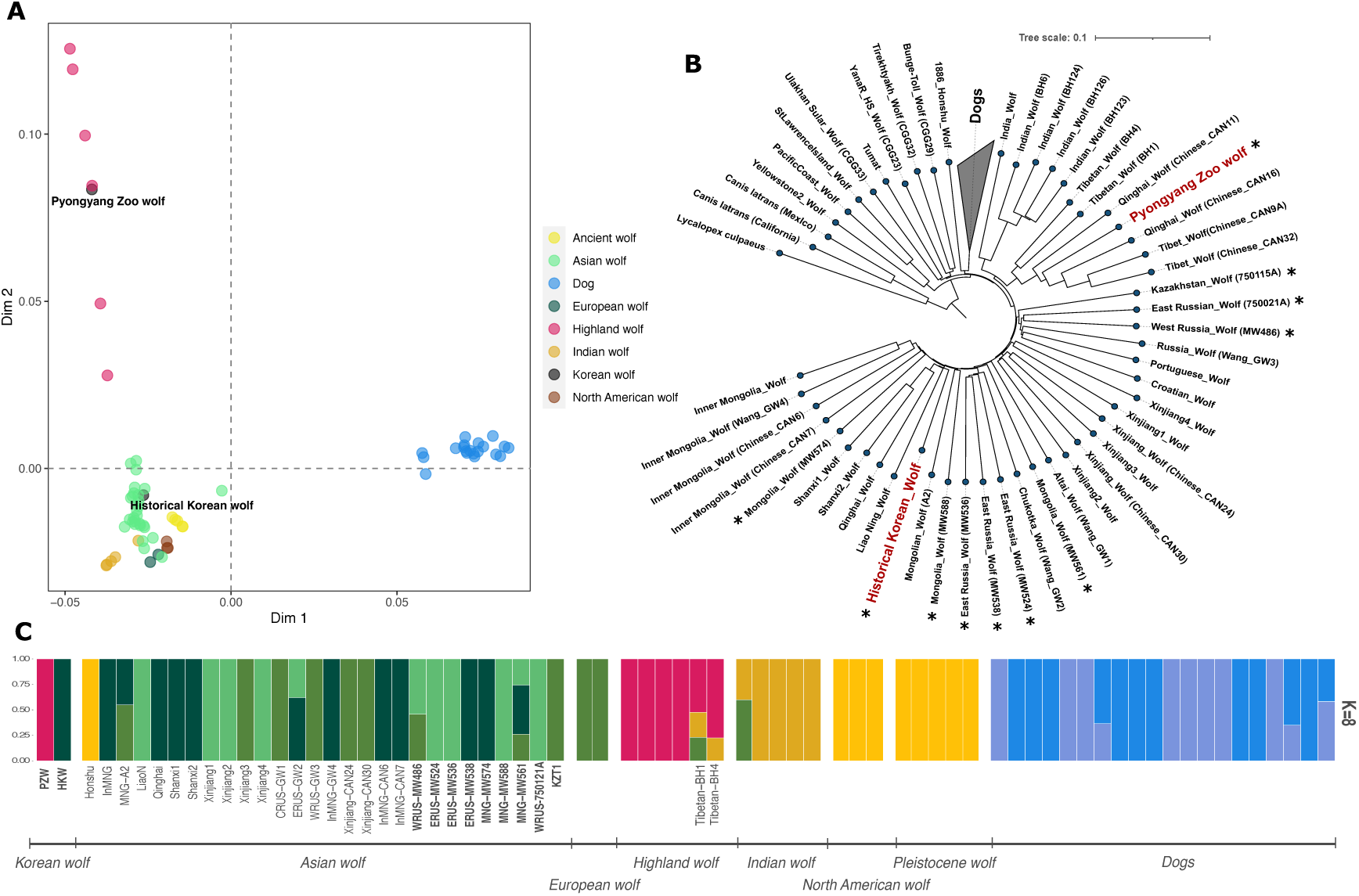
Korean wolf genomic affinities and population structure. A) Multidimensional scaling (MDS) plot using genome-wide data and including all dogs and wolves in the dataset. B) Distance-based NJ tree showing the placement of the historical Korean wolf and the captive Pyongyang Zoo wolf. C) Admixture plot assuming 8 ancestry components (K) showing the historical Korean wolf and the captive Pyongyang Zoo wolf cluster with different groups. Each bar corresponds to a different genome. The colors indicate the inferred ancestry components and their proportions. Asterisk next to sample’s names in the NJ tree as well as bold names in the admixture plot, indicate the novo resequenced wolf genomes.

To elucidate in more detail the internal population structure among the wolves, we performed a second MDS analysis excluding all dog samples. In this case, dimension 1 separates highland wolves from the rest of the wolf samples and dimension 2 separates Indian wolves (Figure S2). The third group of wolves observed in the upper right corner of the plot includes the North American, ancient Siberian, European and Asian wolves. The internal structure of this last-mentioned group of wolves resembles the geographical distribution of the Eurasian samples from east to west separated along dimension 2. Consistent with our previous observation, we here again observe that the PZW and the HKW samples show affinities to different populations, specifically, the PZW is placed close to the Qinghai-CAN11 wolf from central China, and the HKW close to Mongolian and north Chinese wolves. In contrast, the other ten resequenced wolf genomes are placed accordingly with their geographic origin. East Russian wolves (MW536, MW538, and MW524) cluster together with other wolves from the same region, as well as the Mongolian wolves (MW561, MW588, MW574, and MGN-A2) cluster with other Mongolian or Inner Mongolian wolves. Wolves from west Russia (MW486) and west Kazakhstan (KZT1) cluster near to European wolves, except for the west Russian wolf 750021A which appeared between the European wolves cluster and west Chinese, central Russian and some ancient Siberian wolves.

We then performed a clustering analysis using ADMIXTURE (Alexander et al. 2009) to estimate individual ancestries assuming two to ten ancestry components (K). When eight ancestry components were estimated, the recovered clusters included two different groups of dogs, ancient wolves and North American wolves in the same group, Indian wolves, highland wolves, and finally, three groups among Eurasian wolves populations. The results again reveal that the two Korean samples derive from clearly distinct sources, with the HKW clustering principally with wolves from China while the PZW clusters with the highland wolves (Figure 1C). Across the different estimated ancestry components the same pattern can be observed where the PZW clusters with wolves from Tibet and Qinghai, and the HKW clusters with east Asian wolves, mainly from Mongolia and northern China. Like in the MDS results, the rest of the wolves show a clustering pattern that resembles their geographic distribution, including the Asian resequenced wolf genomes (Figure S3).

Similar relationships were recovered from a distance-based Neighbor-Joining (NJ) tree and the maximum likelihood phylogeny built on 1000 concatenated trees (Figure 1B and S4). In both analyses, we recover the same clades, although the relationships of the basal branches differ between both trees. The main clades recovered in the NJ tree correspond to dogs, North American wolves, Pleistocene wolves, and Eurasian wolves. Wolves are further divided into Indian wolves, highland wolves (including PZW), European and west-central Asian wolves, west Chinese wolves (Xinjiang), east Russian and Mongolian wolves, and finally, east Chinese wolves including HKW as basal to wolves from Liao Ning, Qinghai and Shanxi (Figure 1B). The maximum likelihood phylogeny recovered similar groupings. In both phylogenetic reconstructions, the PZW is placed close to highland wolves from Qinghai with a high bootstrap support, while the HKW is placed together with the Liao Ning wolf, which is the geographically closest population to the Korean Peninsula, and it is closely related to other east Chinese wolves (Figure S4).

Altogether, these results suggest that the PZW is not originally from the Korean Peninsula, instead it is closely related to highland wolves from the Tibetan Plateau. We hypothesis that this specimen was obtained by the Pyongyang Central Zoo via some other route (e.g. exchange with another zoo). Unfortunately, the information about this sample is limited. Given the HKW likely represents the original Korean peninsula population and in light of the original questions of this study, for the remaining analyses we focused mainly on this specimen.

### Dog admixture patterns in the historical Korean wolf

Next, we used outgroup *f*_*3*_*-*statistics to explore more deeply the identified relationships, by evaluating the amount of shared genetic drift between HKW, PZW and other wolf populations. The *f*_*3*_*-*statistics results for the PZW were in agreement with the clustering and phylogenetic analyses, showing high shared genetic drift with highland wolves from the Tibetan Plateau (Figure S5). The *f*_*3*_*-*statistics results for the HKW showed the highest shared genetic drift with east Asian wolves, mainly from central and east China, but also HKW with the highland wolves BH1 and BH4 as well as with Shar Pei dog, suggesting gene flow and admixture among these populations (Figure 2A and 2B). We then used TreeMix (Pickrell and Pritchard 2012) to look into potential admixture patterns suggested by the outgroup *f*_*3*_*-*statistics results. This analysis was performed using a subset of samples which includes the HKW and the closest Asian wolves as observed in the outgroup *f*_*3*_*-*statistics and NJ tree, as well as wolves representing relevant clades on the tree (Portuguese, Xinjiang-CAN30, Indian-BH6, Tibet-CAN9A and the PZW), and Asian dog breeds. The overall tree topology recovered is in agreement with the NJ and maximum-likelihood trees, and the migration events estimated are in agreement with the admixture patterns observed before (Figure 2C, Figure S6 and Figure S7). When allowing one to three migration events, we find the Tibetan wolves BH1 and BH4 placed in the same clade as the Indian wolves, but appear to have had gene flow from the HKW lineage and highland wolves, confirming its admixed nature (Figure S6 and Figure S7). The high level of admixture of these two Tibetan wolf samples from Ladakh was already mentioned in the original study where they were described (*cf*. Hennelly et al. 2021). Finally, the treemix admixture graph shows an admixture event between HKW and Shar Pei dog when allowing for an additional migration edge, supporting our outgroup *f*_*3*_*-*statistics results (Figure 2C, Figure S6 and Figure S7).

**Figure 2.**
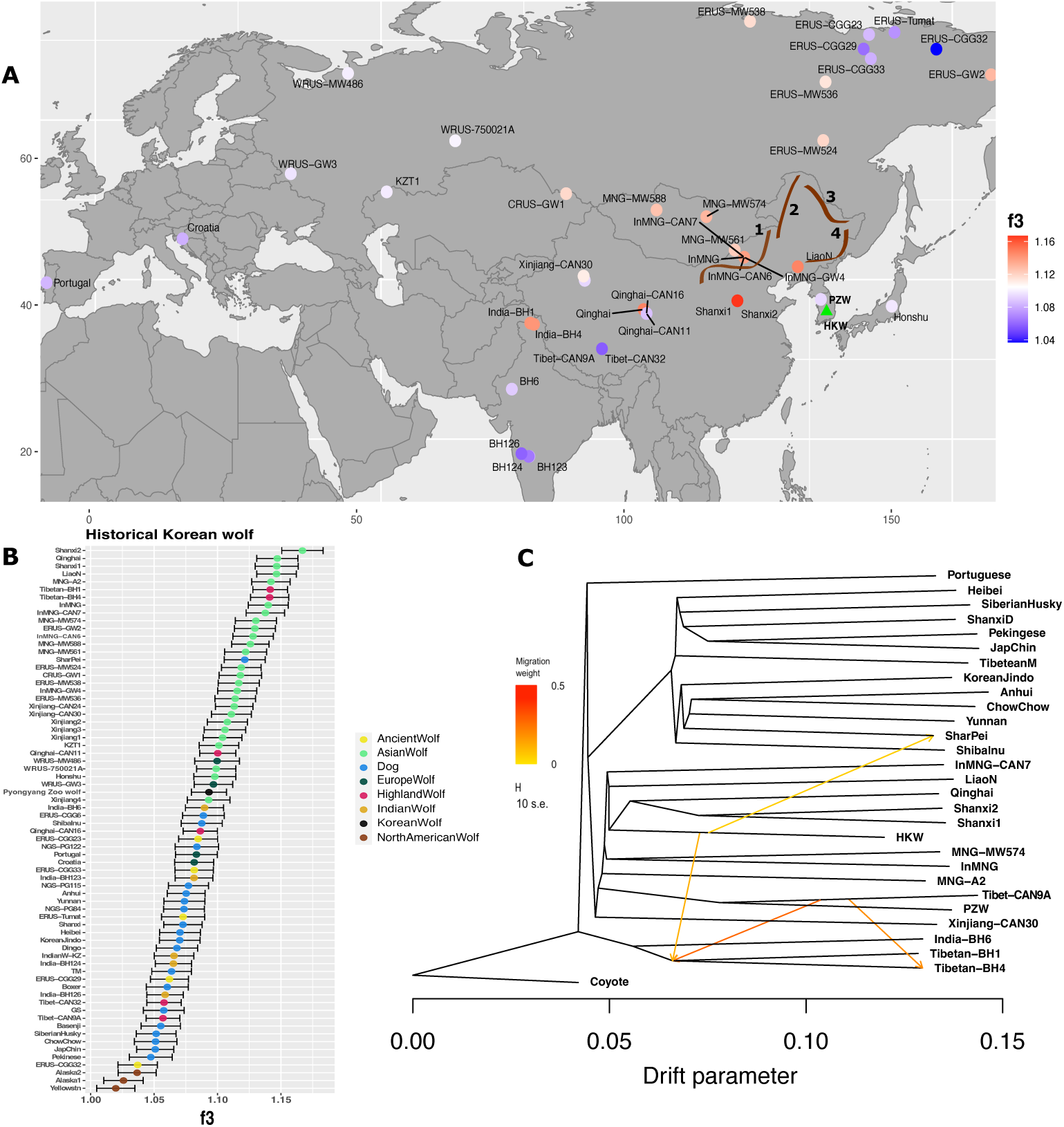
Genetic relationships and admixture patterns estimated by allele frequency data. A) Outgroup *f3-statistics* measuring shared genetic drift between the HKW and Eurasian wolves used in this study (*f3*(HKW, Eurasian wolves; coyote)). Samples are shown in their approximate geographical locations. The samples with highest *f3-statistics* values are shown in red and the lowest in blue. HKW is represented in the map with a green triangle. The drawn lines in the map represent the main geological elevation in the region north to the Korean Peninsula: 1) approximate location of the southeast border of the Mongolian Plateau, 2) the Great Khingan mountains, 3) the Lesser Khingan mountains, and 4) the Changbai mountains. Samples from zoo origin were excluded with the exception of the PZW. The coyotes were used as an outgroup. B) Outgroup *f3-statistics* showing levels of shared genetic drift between the HKW and wolves/dogs. Horizontal lines indicate 3 SE estimated using a block-jackknife procedure in 1Mb blocks. C) TreeMix admixture graph showing the result for 4 migration edges represented by arrows. The arrows indicate admixture events and their color correlates with the intensity of the estimated gene flow between lineages. The analyses were performed on a SNP panel of 1,904,538 transversion sites.

To formally test for admixture between the HKW and dogs, we used *D-*statistics. Firstly, to elucidate the directionality of the gene flow and test the possibility of wolf introgression in the Shar Pei breed, we implemented a test in the form *D*(Andean fox, Eurasian wolves; Tibetan mastiff, Shar Pei). The results showed that Shar Pei dog shares more alleles than the Tibetan mastiff with most of the Eurasian wolves in the test, indicating that Shar Pei dog carries wolf admixture. It is noteworthy the especially high value obtained by HKW in the test when compared with other Eurasian wolf lineages (Figure 3A).

**Figure 3.**
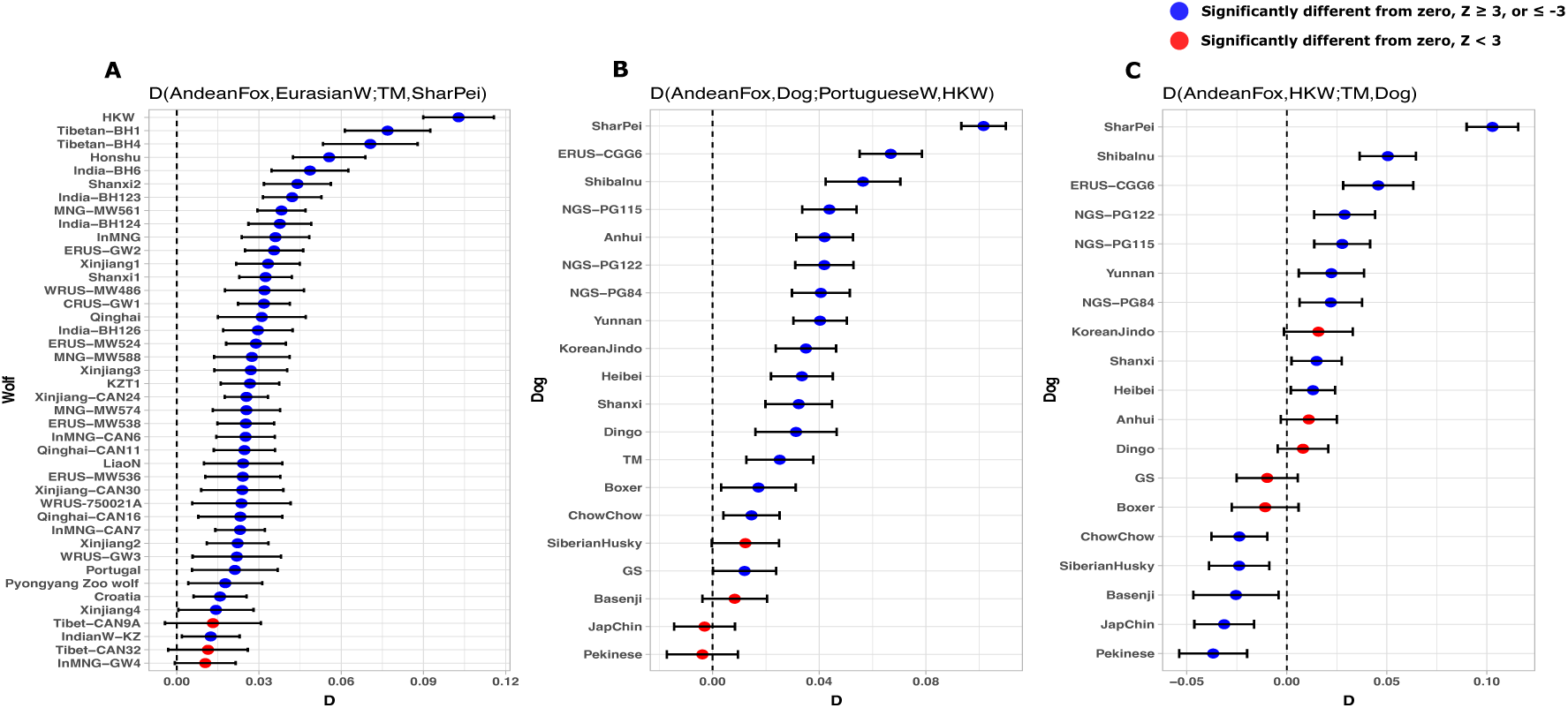
Gene flow between wolves and dog breeds. A) *D-statistics* testing for admixture between wolves and Shar Pei dogs. B-C) *D-statistics* tests showing gene flow between the historical Korean wolf and different dog breeds. In all analyses Andean fox was used as an outgroup, and Tibetan mastiff breed (TM) was used as a representative for dogs. In all cases, Shar Pei appears with the highest *D* values indicating admixture with wolves. Historical Korean wolf shows significant admixture with dogs, in particular with the Shar Pei lineage.

Next, we tested whether the HKW also had admixture from the Shar Pei dog by computing a test of the form *D*(Andean fox, Dog; Portuguese wolf, HKW) (Figure 3B). If HKW carries admixture from the Shar Pei dog, we expect all dogs sharing significantly more alleles with HKW in comparison to the Portuguese wolf. Conversely, if only the Shar Pei dog shares significantly more alleles with HKW, that would imply the gene flow goes only from the HKW into the Shar Pei dog. Similarly, a second test in the form *D*(Andean fox, HKW, Tibetan mastiff, Dog) was performed to explore the dog admixture with the HKW (Figure 3C). In this case, if the HKW had admixed with Shar Pei, we expect most of the dogs sharing significantly more alleles with the HKW, but if Shar Pei dog is admixed with Korean wolves, just the Shar Pei dog should share significantly more alleles with the HKW in comparison to the Tibetan mastiff. The first test confirmed that the HKW is admixed with dogs, and once again, the highest value in the analysis was obtained by the Shar Pei dog, suggesting that this breed is the main genetic source of the HKW admixture with dogs (Figure 3B). The result of the second test supported the admixture patterns observed before, being the Shar Pei dog is the breed with the highest obtained value (Figure 3C). Overall, *D-*statistics and treemix graph results suggest there has been bi-directional gene flow between the HKW to Shar Pei lineages.

### Korean wolf’s genetic diversity

Given the decline of Korean wolf populations and the newly obtained evidence of admixture with the Shar Pei dog, we measured the genetic diversity of the HKW aiming to find clues that support the demographic changes that affected these populations. We estimated the heterozygosity in non-overlapping windows of 1Mb across the first 704 scaffolds longer than 1Mb, using a subset of relevant wolf samples including the HKW and the PZW. When comparing the heterozygosity it is clear that, on average, HKW presented similar levels of heterozygosity than the rest of the wolves, even though a slight tendency to lower heterozygous regions was observed (Figure 4A). Complementary to this analysis, we estimated the inbreeding coefficient (F) of all the modern wolves in our dataset. Our results show low inbreeding levels (0.12) for the HKW compared to other wolves (Figure 4B).

**Figure 4.**
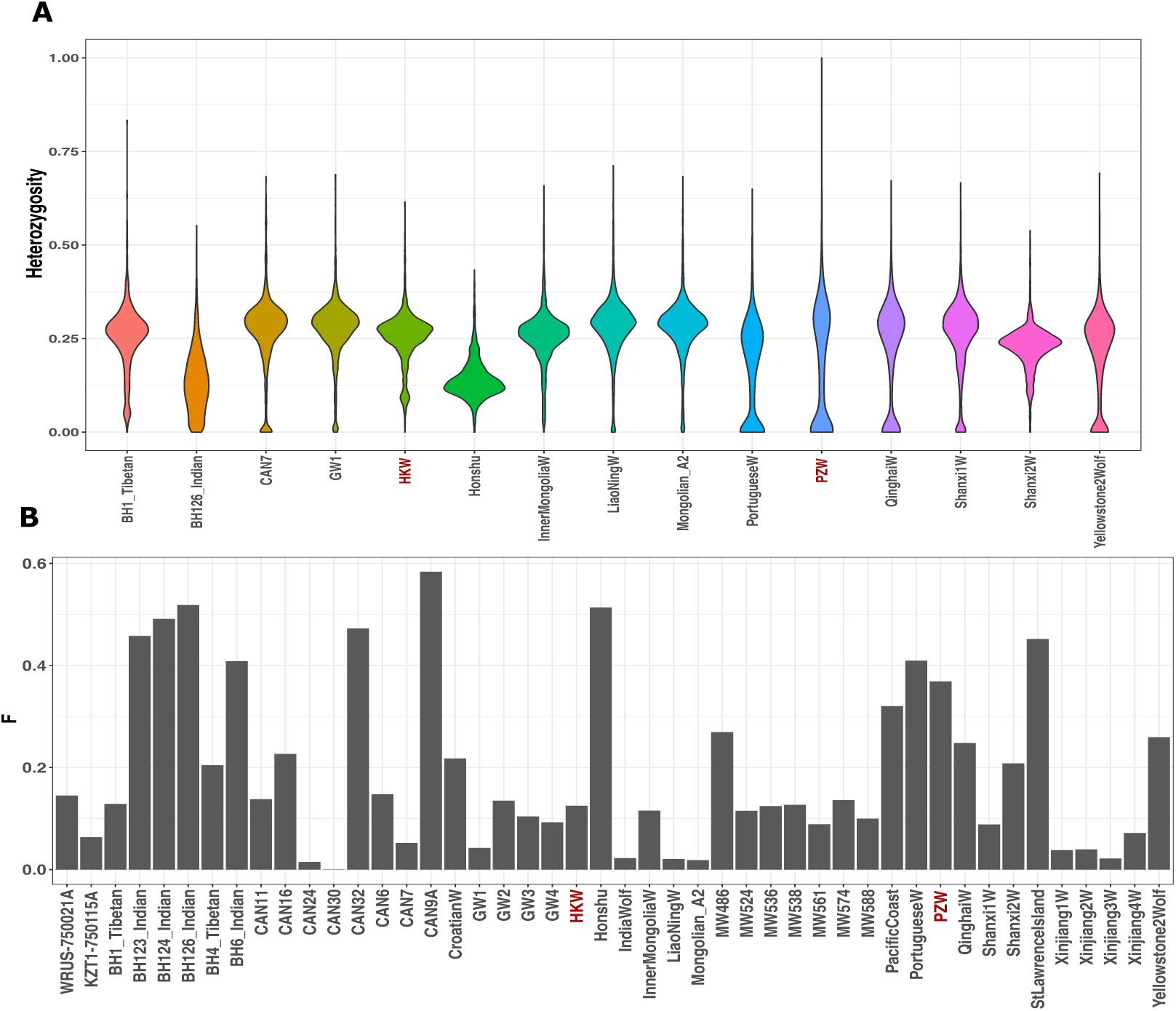
Assessing HKW genetic diversity. A) Heterozygosity estimated in 1Mb genomic windows for representative wolves including the HKW and related samples. The HKW displays similar heterozygosity to most of the compared wolf individuals, however it shows a proportion of low heterozygosity regions. B) Inbreeding coefficient (F) estimated for all modern and historical wolves in our dataset. HKW has a low inbreeding value (0.12) when compared to the rest of wolf samples. Both analyses were performed on genotype imputed data.

## Discussion

Our results demonstrate that Korean wolves are closely related to east Chinese wolf populations and do not seem to be highly genetically differentiated. This implies that wolf populations from the Korean Peninsula were not geographically isolated from the rest of the continental populations as was suggested by Abe, who argued that the Yalou river could serve as a geographical barrier (Abe 1936). Previously published data by Ishiguro et al. (2009; 2010) indicated similar results when analysing the mitochondrial control region. For this reason, and despite the limitation of having used just one wolf sample originally from Korea in this study, we suggest that the observed association patterns correctly reflect the genetic relationships of the likely vanished Korean wolf populations. Therefore, from a genomic perspective, there is no evidence to separate Korean wolves from other closely related populations from China, Mongolia, and Russia currently belonging to the *Canis lupus lupus* subspecies.

Surprisingly, in some analyses, we observed a higher affinity between the HKW and wolves from as far as Qinghai or Ladakh than to wolves from Mongolia or Inner Mongolia that are geographically closer. Our results show regional population structure among the East Asian wolves, where the populations from Inner Mongolia, Mongolia, and east Russia form a group, and the wolves from Korea, Shanxi, Liao Ning, and Qinghai form a second group. Furthermore, the gene flow observed between the HKW and highland wolf populations from Ladakh in the north of India emphasizes such population structure. This suggests that wolf mobility and gene flow between central China or even western Asian regions and eastern China and the Korean Peninsula is more plausible than the mobility to northern regions. Considering the high dispersion capacity of wolves, in the case of the Korean populations, the apparently reduced gene flow with wolves from Inner Mongolia and Mongolia, or from east Russia, probably is not related to topographic barriers like the Mongolian plateau, the Great Khingan Mountains, the Lesser Khingan Mountains, or the Changbai Mountains, but to ecological affinities. It has been observed before the existence of significant isolation by distance among neighboring wolf populations (Geffen et al. 2004) which has been associated with ecological variables such as vegetation type, temperature, or even snow cover, among others (Pilot et al. 2006; Geffen et al. 2004). These habitat preferences have been explained in relationship to prey specialization, due to some of these environmental factors defining the ungulate communities which are the wolves’ main prey (Musiani et al. 2007; Pilot et al. 2006; Carmichael et al. 2001; David Mech and Boitani 2003). Similarly, a north-south differentiation pattern among wolf populations has been described before, again, associated with changes in environmental variables (Stronen et al. 2013; Randi et al. 2001). Then, the grasslands of the Mongolian plateau, or the denser coniferous forests of Russia, as well as the colder weather and other environmental variables associated with those ecosystems could represent less familiar or suitable habitats for Korean wolf populations, being preferable for them to disperse through the North China Plain to the west. In order to corroborate this possible gene flow corridor, it will be important to increase the number of Chinese wolf genomes from regions that are not yet well represented.

Regarding the PZW, this zoo individual clearly seems to be a highland wolf due to its strong affinity with other highland wolves from the Tibetan plateau. The origin of this wolf is uncertain to us. It is possible that the PZW was considered equivalent to the Korean wolf from a taxonomic point of view since, historically, Korean wolves were classified as the subspecies *C. l. chanco*, the same subspecies that has been used to describe highland wolves from the Tibetan plateau. If that is the case, our results suggest Korean wolf conservation programs might not benefit from including the PZW.

Our results are in line with previous studies showing isolated and diminished wolf populations have a higher chance of hybridizing with closely related species, especially dogs (Ciucani et al. 2022; vonHoldt et al. 2022; Niemann et al. 2021; Randi 2001; Phillips et al. 2003; Ballard et al. 2003). Our data revealed that the HKW is admixed with dogs, with the Shar Pei breed being the most likely source of ancestry contributing to the genetic composition of this specimen. The relatively high estimated heterozygosity levels across the genome of the HKW as well as its low inbreeding coefficient estimation could be explained by the admixture with dogs. We could expect to find low heterozygosity levels and inbreeding in populations in decline, as can be observed in the case of the Honshu wolf from Japan, which suffered a similar history of decrease in population size and subsequent extinction (Niemann et al. 2021; Matsumura et al. 2014; Ishiguro et al. 2009). Strong admixture signatures are more common in small and isolated populations, which could be the situation for Korean wolves. However, there is a possibility that the admixture pattern found here represents a particular case of a wolf individual with higher or more recent dog admixture than the average in the Korean wolf populations. To truly elucidate the extent and frequency of dog admixture in the wolf populations from the Korean Peninsula it is necessary to expand the analysis to a larger number of historical specimens from across this region.

In the same way, our results indicate that the Chinese Shar Pei breed is admixed with wolves, mainly from Asia. Taking into consideration that Shar Pei was close to extinction during the 20th century due to its prohibition after the Chinese Communist Revolution (The Editors of Encyclopedia Britannic, 2019), this breed likely went through a population bottleneck, reducing its genetic diversity and favoring admixture with wolves. According to historical records about the Shar Pei dog, the breed survived in Taiwan and Hong Kong, making it unlikely that the main wolf source of admixture were wolves from Korea. Therefore, we consider that the high affinity between the HKW and the Shar Pei dog observed in the *D-* statistics, is due to the Shar Pei ancestry in the Koren wolf rather than the other way around.

Wolves are apex predators that play an important role in the ecosystems they inhabit. The local extirpation of wolves have severe cascading consequences for the trophic chain, unbalancing the distributions and interactions among predators, herbivores and primary producers, as have been well documented in the wolves reintroduction project in the Yellowstone National Park, in the USA (Ripple and Beschta 2012). For this reason, maintaining healthy wolf populations or ensuring a successful reintroduction program could be important to maintain the proper functioning of ecosystems.

Our findings have the potential to contribute to wolf conservation strategies, both at a local level for the reintroduction of wolves in the Korean Peninsula and at a regional level. In the context of the Korean Peninsula, clarifying the genetic affinities of Korean wolves could allow to define an appropriate genetic pool for wolf populations used in conservation and reintroduction programs. Our preliminary results suggest wolves from neighboring regions to the Korean Peninsula, such as northern China would be the best option for replacing the extirpated Korean populations. Perhaps less desirable could be the use of populations from the Mongolian Plateau or eastern Russia for this purpose; nevertheless, further historical samples from Korean wolves could be necessary to confirm it. At a regional level, the detection, protection or creation of biological corridors that could allow the dispersion and gene flow of wolf populations in central and eastern China, connecting with the Korean Peninsula, would be ideal for the protection of the species. This would have the potential to also benefit other threatened species including prey species.

## Acknowledgments

This work was supported by ERC Consolidator Award 681396 Extinction Genomics, DNRF143 Center for Evolutionary Hologenomics, and the Norwegian Environment Agency (project 18088069). G.H-A. is supported by the Consejo Nacional de Ciencia y Tecnología from Mexico (CONACyT, 576743). Morten Skage, Mikeal Åkersson, Jouni Aspi, Kjetill S. Jakobsen, and Øyvind Øverli provided some of the wolf samples included in this study. We would like to acknowledge Zoo Zürich for providing the study with a sample from their Mongolian wolf.

## Data Accessibility and Benefit-Sharing

### Data Accessibility

Generated raw sequence reads are deposited to the European Nucleotide Archive (Project ID: PRJEB57591).

### Benefit-Sharing Statement

The research work presented here was the result of an international collaboration that involved the participation and co-authorship of scientists from the different countries providing the wolf samples. The results produced by this collaboration can directly impact on the local and regional conservation strategies of wolf populations.

## Author contributions

G.H-A. designed the study. K.Y.E., S.C., B.B., G.G., Z.U., and P.A.K. provided the wolf samples. C.H.S-O., M-H.S.S., L.T.L, C.G.C., M.M.C, S.S.T.M., and N.F.M. did the laboratory processing of the samples. J.B., S.J., and C.K. performed the whole-genome resequencing procedure of the HKW sample. G.H-A. performed the data analysis with support from J.R-M., and S.G. X.S. performed the genotype imputation. G.H-A wrote the manuscript with support from J.R-M., S.G., and M.T.P.G. N.F.M and M-H.S.S wrote the methods related to the laboratory processing of the samples. Funding was obtained by M.T.P.G. and H.K.S. All authors revised the final manuscript.

## References

Abe, Y. 1923. On the Korean wolf. Zoological Magazine, 35, 380–386.

Abe, Y. 1936. On the Corean wolf again (an Answer to Mr. Pocock). Zoological Magazine, 48, 639–644.

Aggarwal, R.K., Kivisild, T., Ramadevi, J., Singh, L., 2007. Mitochondrial DNA coding region sequences support the phylogenetic distinction of two Indian wolf species. Journal of Zoological Systematics and Evolutionary Research 45, 163–172. https://doi.org/10.1111/j.1439-0469.2006.00400.x

Aggarwal, R.K., Ramadevi, J., Singh, L., 2003. Ancient origin and evolution of the Indian wolf: evidence from mitochondrial DNA typing of wolves from Trans-Himalayan region and Peninsular India. Genome Biology 4, P6. https://doi.org/10.1186/gb-2003-4-6-p6

Alexander, D.H., Novembre, J., Lange, K., 2009. Fast model-based estimation of ancestry in unrelated individuals. Genome Res. 19, 1655–1664. https://doi.org/10.1101/gr.094052.109

Alvares, F., Bogdanowicz, W., Campbell, L.A.D., Godinho, R., Jhala, Y.V., Kitchener, A.C., Koepfli, K.-P., Krofel, M., Sillero-Zubiri, C., Viranta, S., Werhahn, G., n.d. Old World Canis spp. with taxonomic ambiguity: Workshop conclusions and recommendations 8.

Auton, A., Li, Y.R., Kidd, J., Oliveira, K., Nadel, J., Holloway, J.K., Hayward, J.J., Cohen, P.E., Greally, J.M., Wang, J., Bustamante, C.D., Boyko, A.R., 2013. Genetic Recombination Is Targeted towards Gene Promoter Regions in Dogs. PLOS Genetics 9, e1003984. https://doi.org/10.1371/journal.pgen.1003984

Ballard, W., Carbyn, L., Smith, D., 2003. Wolf Interactions with Non-prey. USGS Northern Prairie Wildlife Research Center.

Encyclopedia Britannica. Chinese shar-pei. https://www.britannica.com/animal/Chinese-shar-pei.

Boessenkool, S., Hanghøj, K., Nistelberger, H.M., Der Sarkissian, C., Gondek, A.T., Orlando, L., Barrett, J.H., Star, B., 2017. Combining bleach and mild predigestion improves ancient DNA recovery from bones. Molecular Ecology Resources 17, 742–751. https://doi.org/10.1111/1755-0998.12623

Browning, B.L., Zhou, Y., Browning, S.R., 2018. A One-Penny Imputed Genome from Next-Generation Reference Panels. The American Journal of Human Genetics 103, 338–348. https://doi.org/10.1016/j.ajhg.2018.07.015

Carmichael, L.E., Nagy, J.A., Larter, N.C., Strobeck, C., 2001. Prey specialization may influence patterns of gene flow in wolves of the Canadian Northwest. Molecular Ecology 10, 2787–2798. https://doi.org/10.1046/j.0962-1083.2001.01408.x

Carøe, C., Gopalakrishnan, S., Vinner, L., Mak, S.S.T., Sinding, M.H.S., Samaniego, J.A., Wales, N., Sicheritz-Pontén, T., Gilbert, M.T.P., 2018. Single-tube library preparation for degraded DNA. Methods in Ecology and Evolution 9, 410–419. https://doi.org/10.1111/2041-210X.12871

Chang, C.C., Chow, C.C., Tellier, L.C., Vattikuti, S., Purcell, S.M., Lee, J.J., 2015. Second-generation PLINK: rising to the challenge of larger and richer datasets. GigaScience 4, s13742-015-0047–8. https://doi.org/10.1186/s13742-015-0047-8

Ciucani, M.M., Ramos-Madrigal, J., Hernández-Alonso, G., Carmagnini, A., Aninta, S.G., Scharff-Olsen, C.H., Lanigan, L.T., Fracasso, I., Clausen, C.G., Aspi, J., Kojola, I., Baltrunaite, L., Balciauskas, L., Moore, J., Åkesson, M., Saarma, U., Hindrikson, M., Hulva, P., Bolfíková, B.C., Nowak, C., Godinho, R., Smith, S., Paule, L., Nowak, S., Myslajek, R.W., Brutto, S.L., Ciucci, P., Boitani, L., Vernesi, C., Stenøien, H.K., Smith, O., Frantz, L., Rossi, L., Angelici, F.M., Cilli, E., Sinding, M.-H.S., Gilbert, M.T.P., Gopalakrishnan, S., 2022. Genomes of the extinct Sicilian wolf reveal a complex history of isolation and admixture with ancient dogs. https://doi.org/10.1101/2022.01.21.477289

Dabney, J., Knapp, M., Glocke, I., Gansauge, M.-T., Weihmann, A., Nickel, B., Valdiosera, C., García, N., Pääbo, S., Arsuaga, J.-L., Meyer, M., 2013. Complete mitochondrial genome sequence of a Middle Pleistocene cave bear reconstructed from ultrashort DNA fragments. Proc. Natl. Acad. Sci. U.S.A. 110, 15758–15763. https://doi.org/10.1073/pnas.1314445110

Danecek, P., Auton, A., Abecasis, G., Albers, C.A., Banks, E., DePristo, M.A., Handsaker, R.E., Lunter, G., Marth, G.T., Sherry, S.T., McVean, G., Durbin, R., 1000 Genomes Project Analysis Groupxs, 2011. The variant call format and VCFtools. Bioinformatics 27, 2156–2158. https://doi.org/10.1093/bioinformatics/btr330

Danecek, P., McCarthy, S.A., 2017. BCFtools/csq: haplotype-aware variant consequences. Bioinformatics 33, 2037–2039. https://doi.org/10.1093/bioinformatics/btx100

Decker, B., Davis, B.W., Rimbault, M., Long, A.H., Karlins, E., Jagannathan, V., Reiman, R., Parker, H.G., Drögemüller, C., Corneveaux, J.J., Chapman, E.S., Trent, J.M., Leeb, T., Huentelman, M.J., Wayne, R.K., Karyadi, D.M., Ostrander, E.A., 2015. Comparison against 186 canid whole-genome sequences reveals survival strategies of an ancient clonally transmissible canine tumor. Genome Res. 25, 1646–1655. https://doi.org/10.1101/gr.190314.115

DePristo, M.A., Banks, E., Poplin, R., Garimella, K.V., Maguire, J.R., Hartl, C., Philippakis, A.A., del Angel, G., Rivas, M.A., Hanna, M., McKenna, A., Fennell, T.J., Kernytsky, A.M., Sivachenko, A.Y., Cibulskis, K., Gabriel, S.B., Altshuler, D., Daly, M.J., 2011. A framework for variation discovery and genotyping using next-generation DNA sequencing data. Nat Genet 43, 491–498. https://doi.org/10.1038/ng.806

Fan, Z., Silva, P., Gronau, I., Wang, S., Armero, A.S., Schweizer, R.M., Ramirez, O., Pollinger, J., Galaverni, M., Del-Vecchyo, D.O., Du, L., Zhang, W., Zhang, Z., Xing, J., Vilà, C., Marques-Bonet, T., Godinho, R., Yue, B., Wayne, R.K., 2016. Worldwide patterns of genomic variation and admixture in gray wolves. Genome Res. 26, 163–173. https://doi.org/10.1101/gr.197517.115

Francis, R.M., 2017. pophelper: an R package and web app to analyse and visualize population structure. Molecular Ecology Resources 17, 27–32. https://doi.org/10.1111/1755-0998.12509

Freedman, A.H., Gronau, I., Schweizer, R.M., Vecchyo, D.O.-D., Han, E., Silva, P.M., Galaverni, M., Fan, Z., Marx, P., Lorente-Galdos, B., Beale, H., Ramirez, O., Hormozdiari, F., Alkan, C., Vilà, C., Squire, K., Geffen, E., Kusak, J., Boyko, A.R., Parker, H.G., Lee, C., Tadigotla, V., Siepel, A., Bustamante, C.D., Harkins, T.T., Nelson, S.F., Ostrander, E.A., Marques-Bonet, T., Wayne, R.K., Novembre, J., 2014. Genome Sequencing Highlights the Dynamic Early History of Dogs. PLOS Genetics 10, e1004016. https://doi.org/10.1371/journal.pgen.1004016

Geffen, E., Anderson, M.J., Wayne, R.K., 2004. Climate and habitat barriers to dispersal in the highly mobile grey wolf. Molecular Ecology 13, 2481–2490. https://doi.org/10.1111/j.1365-294X.2004.02244.x

Gilbert, M.T.P., Tomsho, L.P., Rendulic, S., Packard, M., Drautz, D.I., Sher, A., Tikhonov, A., Dalén, L., Kuznetsova, T., Kosintsev, P., Campos, P.F., Higham, T., Collins, M.J., Wilson, A.S., Shidlovskiy, F., Buigues, B., Ericson, P.G.P., Germonpré, M., Götherström, A., Iacumin, P., Nikolaev, V., Nowak-Kemp, M., Willerslev, E., Knight, J.R., Irzyk, G.P., Perbost, C.S., Fredrikson, K.M., Harkins, T.T., Sheridan, S., Miller, W., Schuster, S.C., 2007. Whole-Genome Shotgun Sequencing of Mitochondria from Ancient Hair Shafts. Science 317, 1927–1930. https://doi.org/10.1126/science.1146971

Gopalakrishnan, S., Samaniego Castruita, J.A., Sinding, M.-H.S., Kuderna, L.F.K., Räikkönen, J., Petersen, B., Sicheritz-Ponten, T., Larson, G., Orlando, L., Marques-Bonet, T., Hansen, A.J., Dalén, L., Gilbert, M.T.P., 2017. The wolf reference genome sequence (Canis lupus lupus) and its implications for Canis spp. population genomics. BMC Genomics 18, 495. https://doi.org/10.1186/s12864-017-3883-3

Gopalakrishnan, S., Sinding, M.-H.S., Ramos-Madrigal, J., Niemann, J., Samaniego Castruita, J.A., Vieira, F.G., Carøe, C., Montero, M. de M., Kuderna, L., Serres, A., González-Basallote, V.M., Liu, Y.-H., Wang, G.-D., Marques-Bonet, T., Mirarab, S., Fernandes, C., Gaubert, P., Koepfli, K.-P., Budd, J., Rueness, E.K., Sillero, C., Heide-Jørgensen, M.P., Petersen, B., Sicheritz-Ponten, T., Bachmann, L., Wiig, Ø., Hansen, A.J., Gilbert, M.T.P., 2018. Interspecific Gene Flow Shaped the Evolution of the Genus Canis. Current Biology 28, 3441-3449.e5. https://doi.org/10.1016/j.cub.2018.08.041

Gray, J. E. 1863. Notice of the chanco or golden wolf (Canis chanco) from Chinsese Tartary. Proceedings of the Zoological Society of London, 31, 94.

Hennelly, L.M., Habib, B., Modi, S., Rueness, E.K., Gaubert, P., Sacks, B.N., 2021. Ancient divergence of Indian and Tibetan wolves revealed by recombination-aware phylogenomics. Molecular Ecology 30, 6687–6700. https://doi.org/10.1111/mec.16127

Ishiguro, N., Inoshima, Y., Shigehara, N., 2009. Mitochondrial DNA Analysis of the Japanese Wolf (Canis Lupus Hodophilax Temminck, 1839) and Comparison with Representative Wolf and Domestic Dog Haplotypes. jzoo 26, 765–770. https://doi.org/10.2108/zsj.26.765

Ishiguro, N., Inoshima, Y., Shigehara, N., Ichikawa, H., Kato, M., 2010. Osteological and Genetic Analysis of the Extinct Ezo Wolf (Canis Lupus Hattai) from Hokkaido Island, Japan. jzoo 27, 320–324. https://doi.org/10.2108/zsj.27.320

Jo, Y.-S., Baccus, J.T., Koprowski, J.L., 2018. Mammals of Korea: a review of their taxonomy, distribution and conservation status. Zootaxa 4522, 1–216. https://doi.org/10.11646/zootaxa.4522.1.1

Jónsson, H., Ginolhac, A., Schubert, M., Johnson, P.L.F., Orlando, L., 2013. mapDamage2.0: fast approximate Bayesian estimates of ancient DNA damage parameters. Bioinformatics 29, 1682–1684. https://doi.org/10.1093/bioinformatics/btt193

Joshi, B., Lyngdoh, S., Singh, S.K., Sharma, R., Kumar, V., Tiwari, V.P., Dar, S.A., Maheswari, A., Pal, R., Bashir, T., Reshamwala, H.S., Shrotriya, S., Sathyakumar, S., Habib, B., Kvist, L., Goyal, S.P., 2020. Revisiting the Woolly wolf (Canis lupus chanco) phylogeny in Himalaya: Addressing taxonomy, spatial extent and distribution of an ancient lineage in Asia. PLOS ONE 15, e0231621. https://doi.org/10.1371/journal.pone.0231621

Kapp, J.D., Green, R.E., Shapiro, B., 2021. A Fast and Efficient Single-stranded Genomic Library Preparation Method Optimized for Ancient DNA. Journal of Heredity 112, 241–249. https://doi.org/10.1093/jhered/esab012

Kim, R.N., Kim, D.-S., Choi, S.-H., Yoon, B.-H., Kang, A., Nam, S.-H., Kim, D.-W., Kim, J.-J., Ha, J.-H., Toyoda, A., Fujiyama, A., Kim, A., Kim, M.-Y., Park, K.-H., Lee, K.S., Park, H.-S., 2012. Genome Analysis of the Domestic Dog (Korean Jindo) by Massively Parallel Sequencing. DNA Research 19, 275–288. https://doi.org/10.1093/dnares/dss011

Kolicheski, A., Johnson, G. s., Villani, N. a., O’Brien, D. p., Mhlanga-Mutangadura, T., Wenger, D. a., Mikoloski, K., Eagleson, J. s., Taylor, J. f., Schnabel, R. d., Katz, M. l., 2017. GM2 Gangliosidosis in Shiba Inu Dogs with an In-Frame Deletion in HEXB. Journal of Veterinary Internal Medicine 31, 1520–1526. https://doi.org/10.1111/jvim.14794

Korneliussen, T.S., Albrechtsen, A., Nielsen, R., 2014. ANGSD: Analysis of Next Generation Sequencing Data. BMC Bioinformatics 15, 356. https://doi.org/10.1186/s12859-014-0356-4

Kozlov, A.M., Darriba, D., Flouri, T., Morel, B., Stamatakis, A., 2019. RAxML-NG: a fast, scalable and user-friendly tool for maximum likelihood phylogenetic inference. Bioinformatics 35, 4453–4455. https://doi.org/10.1093/bioinformatics/btz305

Letunic, I., Bork, P., 2019. Interactive Tree Of Life (iTOL) v4: recent updates and new developments. Nucleic Acids Research 47, W256–W259. https://doi.org/10.1093/nar/gkz239

Li, H., Durbin, R., 2009. Fast and accurate short read alignment with Burrows–Wheeler transform. Bioinformatics 25, 1754–1760. https://doi.org/10.1093/bioinformatics/btp324

Lindblad-Toh, K., Wade, C.M., Mikkelsen, T.S., Karlsson, E.K., Jaffe, D.B., Kamal, M., Clamp, M., Chang, J.L., Kulbokas, E.J., Zody, M.C., Mauceli, E., Xie, X., Breen, M., Wayne, R.K., Ostrander, E.A., Ponting, C.P., Galibert, F., Smith, D.R., deJong, P.J., Kirkness, E., Alvarez, P., Biagi, T., Brockman, W., Butler, J., Chin, C.-W., Cook, A., Cuff, J., Daly, M.J., DeCaprio, D., Gnerre, S., Grabherr, M., Kellis, M., Kleber, M., Bardeleben, C., Goodstadt, L., Heger, A., Hitte, C., Kim, L., Koepfli, K.-P., Parker, H.G., Pollinger, J.P., Searle, S.M.J., Sutter, N.B., Thomas, R., Webber, C., Lander, E.S., 2005. Genome sequence, comparative analysis and haplotype structure of the domestic dog. Nature 438, 803–819. https://doi.org/10.1038/nature04338

Mak, S.S.T., Gopalakrishnan, S., Carøe, C., Geng, C., Liu, S., Sinding, M.-H.S., Kuderna, L.F.K., Zhang, W., Fu, S., Vieira, F.G., Germonpré, M., Bocherens, H., Fedorov, S., Petersen, B., Sicheritz-Pontén, T., Marques-Bonet, T., Zhang, G., Jiang, H., Gilbert, M.T.P., 2017. Comparative performance of the BGISEQ-500 vs Illumina HiSeq2500 sequencing platforms for palaeogenomic sequencing. GigaScience 6, gix049. https://doi.org/10.1093/gigascience/gix049

Marchant, T.W., Johnson, E.J., McTeir, L., Johnson, C.I., Gow, A., Liuti, T., Kuehn, D., Svenson, K., Bermingham, M.L., Drögemüller, M., Nussbaumer, M., Davey, M.G., Argyle, D.J., Powell, R.M., Guilherme, S., Lang, J., Ter Haar, G., Leeb, T., Schwarz, T., Mellanby, R.J., Clements, D.N., Schoenebeck, J.J., 2017. Canine Brachycephaly Is Associated with a Retrotransposon-Mediated Missplicing of SMOC2. Current Biology 27, 1573-1584.e6. https://doi.org/10.1016/j.cub.2017.04.057

Marsden, C.D., Ortega-Del Vecchyo, D., O’Brien, D.P., Taylor, J.F., Ramirez, O., Vilà, C., Marques-Bonet, T., Schnabel, R.D., Wayne, R.K., Lohmueller, K.E., 2016. Bottlenecks and selective sweeps during domestication have increased deleterious genetic variation in dogs. Proceedings of the National Academy of Sciences 113, 152–157. https://doi.org/10.1073/pnas.1512501113

Matsumura, S., Inoshima, Y., Ishiguro, N., 2014. Reconstructing the colonization history of lost wolf lineages by the analysis of the mitochondrial genome. Molecular Phylogenetics and Evolution 80, 105–112. https://doi.org/10.1016/j.ympev.2014.08.004

McKenna, A., Hanna, M., Banks, E., Sivachenko, A., Cibulskis, K., Kernytsky, A., Garimella, K., Altshuler, D., Gabriel, S., Daly, M., DePristo, M.A., 2010. The Genome Analysis Toolkit: A MapReduce framework for analyzing next-generation DNA sequencing data. Genome Res. 20, 1297–1303. https://doi.org/10.1101/gr.107524.110

Mech, L., Boitani, L., 2003. Wolf Social Ecology. USGS Northern Prairie Wildlife Research Center.

Metzger, J., Nolte, A., Uhde, A.-K., Hewicker-Trautwein, M., Distl, O., 2017. Whole genome sequencing identifies missense mutation in MTBP in Shar-Pei affected with Autoinflammatory Disease (SPAID). BMC Genomics 18, 348. https://doi.org/10.1186/s12864-017-3737-z

Mivart, S. G. 1890. The Common Wolf. In Dogs, jackals, wolves, and foxes: a monograph of the Canidæ. (Tylor and Francis), pp. 8. https://doi.org/10.5962/bhl.title.23888

Musiani, M., Leonard, J.A., Cluff, H.D., Gates, C.C., Mariani, S., Paquet, P.C., Vilà, C., Wayne, R.K., 2007. Differentiation of tundra/taiga and boreal coniferous forest wolves: genetics, coat colour and association with migratory caribou. Molecular Ecology 16, 4149–4170. https://doi.org/10.1111/j.1365-294X.2007.03458.x

National Institute of Biological Resources. 2012. Red Data Book of Endangered Mammals in Korea. National Institute of Biological Resources. pp. 111.

Niemann, J., Gopalakrishnan, S., Yamaguchi, N., Ramos-Madrigal, J., Wales, N., Gilbert, M.T.P., Sinding, M.-H.S., 2021. Extended survival of Pleistocene Siberian wolves into the early 20th century on the island of Honshu. iScience 24, 101904. https://doi.org/10.1016/j.isci.2020.101904

Nowak, R.M., 2010. Wolf Evolution and Taxonomy. In Wolves: behavior, ecology, and conservation, L. D. Mech and L. Boitani, ed.(University of Chicago Press), pp. 239–258. https://doi.org/10.7208/9780226516981-013

Nowak, R. M. 1995. Another look at wolf taxonomy. In Ecology and conservation of wolves in a changing world, L. N. Carbyn, S. H. Fritts and D. R. Seip, ed. (Canadian Circumpolar Institute), pp. 375–398.

Paradis, E., Schliep, K., 2019. ape 5.0: an environment for modern phylogenetics and evolutionary analyses in R. Bioinformatics 35, 526–528. https://doi.org/10.1093/bioinformatics/bty633

Patterson, N., Moorjani, P., Luo, Y., Mallick, S., Rohland, N., Zhan, Y., Genschoreck, T., Webster, T., Reich, D., 2012. Ancient Admixture in Human History. Genetics 192, 1065–1093. https://doi.org/10.1534/genetics.112.145037

Phillips, M., Henry, V., Kelly, B., 2003. Restoration of the Red Wolf. USGS Northern Prairie Wildlife Research Center.

Pickrell, J., Pritchard, J., 2012. Inference of population splits and mixtures from genome-wide allele frequency data. Nat Prec 1–1. https://doi.org/10.1038/npre.2012.6956.1

Pilot, M., Jedrzejewski, W., Branicki, W., Sidorovich, V.E., Jedrzejewska, B., Stachura, K., Funk, S.M., 2006. Ecological factors influence population genetic structure of European grey wolves. Molecular Ecology 15, 4533–4553. https://doi.org/10.1111/j.1365-294X.2006.03110.x

Ramos-Madrigal, J., Sinding, M.-H.S., Carøe, C., Mak, S.S.T., Niemann, J., Samaniego Castruita, J.A., Fedorov, S., Kandyba, A., Germonpré, M., Bocherens, H., Feuerborn, T.R., Pitulko, V.V., Pavlova, E.Y., Nikolskiy, P.A., Kasparov, A.K., Ivanova, V.V., Larson, G., Frantz, L.A.F., Willerslev, E., Meldgaard, M., Petersen, B., Sicheritz-Ponten, T., Bachmann, L., Wiig, Ø., Hansen, A.J., Gilbert, M.T.P., Gopalakrishnan, S., 2021. Genomes of Pleistocene Siberian Wolves Uncover Multiple Extinct Wolf Lineages. Current Biology 31, 198-206.e8. https://doi.org/10.1016/j.cub.2020.10.002

Randi, E., Lucchini, V., Christensen, M.F., Mucci, N., Funk, S.M., Dolf, G., Loeschcke, V., 2001. Mitochondrial DNA Variability in Italian and East European Wolves: Detecting the Consequences of Small Population Size and Hybridization. Conservation Biology 14, 464–473. https://doi.org/10.1046/j.1523-1739.2000.98280.x

Ripple, W.J., Beschta, R.L., 2012. Trophic cascades in Yellowstone: The first 15years after wolf reintroduction. Biological Conservation 145, 205–213. https://doi.org/10.1016/j.biocon.2011.11.005

Schubert, M., Ermini, L., Sarkissian, C.D., Jónsson, H., Ginolhac, A., Schaefer, R., Martin, M.D., Fernández, R., Kircher, M., McCue, M., Willerslev, E., Orlando, L., 2014. Characterization of ancient and modern genomes by SNP detection and phylogenomic and metagenomic analysis using PALEOMIX. Nat Protoc 9, 1056–1082. https://doi.org/10.1038/nprot.2014.063

Schubert, M., Lindgreen, S., Orlando, L., 2016. AdapterRemoval v2: rapid adapter trimming, identification, and read merging. BMC Research Notes 9, 88. https://doi.org/10.1186/s13104-016-1900-2

Shrotriya, S., Lyngdoh, S., Habib, B., 2012. Wolves in Trans-Himalayas: 165 years of taxonomic confusion. Current Science 103, 885–887.

Sinding, M.-H.S., Gopalakrishan, S., Vieira, F.G., Castruita, J.A.S., Raundrup, K., Jørgensen, M.P.H., Meldgaard, M., Petersen, B., Sicheritz-Ponten, T., Mikkelsen, J.B., Marquard-Petersen, U., Dietz, R., Sonne, C., Dalén, L., Bachmann, L., Wiig, Ø., Hansen, A.J., Gilbert, M.T.P., 2018. Population genomics of grey wolves and wolf-like canids in North America. PLOS Genetics 14, e1007745. https://doi.org/10.1371/journal.pgen.1007745

Sinding, M.-H.S., Gopalakrishnan, S., Ramos-Madrigal, J., de Manuel, M., Pitulko, V.V., Kuderna, L., Feuerborn, T.R., Frantz, L.A.F., Vieira, F.G., Niemann, J., Samaniego Castruita, J.A., Carøe, C., Andersen-Ranberg, E.U., Jordan, P.D., Pavlova, E.Y., Nikolskiy, P.A., Kasparov, A.K., Ivanova, V.V., Willerslev, E., Skoglund, P., Fredholm, M., Wennerberg, S.E., Heide-Jørgensen, M.P., Dietz, R., Sonne, C., Meldgaard, M., Dalén, L., Larson, G., Petersen, B., Sicheritz-Pontén, T., Bachmann, L., Wiig, Ø., Marques-Bonet, T., Hansen, A.J., Gilbert, M.T.P., 2020. Arctic-adapted dogs emerged at the Pleistocene–Holocene transition. Science 368, 1495–1499. https://doi.org/10.1126/science.aaz8599

Sokolov, V. E., O. L. Rossolimo. 1985. Taxonomy and variability. In The wolf: History, systematics, morphology and ecology, D. I. Bibikov, ed. (Nauka), pp. 21–50.

Stronen, A.V., Jedrzejewska, B., Pertoldi, C., Demontis, D., Randi, E., Niedzialkowska, M., Pilot, M., Sidorovich, V.E., Dykyy, I., Kusak, J., Tsingarska, E., Kojola, I., Karamanlidis, A.A., Ornicans, A., Lobkov, V.A., Dumenko, V., Czarnomska, S.D., 2013. North-South Differentiation and a Region of High Diversity in European Wolves (Canis lupus). PLOS ONE 8, e76454. https://doi.org/10.1371/journal.pone.0076454

vonHoldt, B.M., Cahill, J.A., Fan, Z., Gronau, I., Robinson, J., Pollinger, J.P., Shapiro, B., Wall, J., Wayne, R.K., 2016. Whole-genome sequence analysis shows that two endemic species of North American wolf are admixtures of the coyote and gray wolf. Science Advances 2, e1501714. https://doi.org/10.1126/sciadv.1501714

vonHoldt, B.M., Hinton, J.W., Shutt, A.C., Murphy, S.M., Karlin, M.L., Adams, J.R., Waits, L.P., Brzeski, K.E., 2022. Reviving ghost alleles: Genetically admixed coyotes along the American Gulf Coast are critical for saving the endangered red wolf. Science Advances 8, eabn7731. https://doi.org/10.1126/sciadv.abn7731

Wang, G., Zhai, W., Yang, H., Fan, R., Cao, X., Zhong, L., Wang, L., Liu, F., Wu, H., Cheng, L., Poyarkov, A.D., Poyarkov JR, N.A., Tang, S., Zhao, W., Gao, Y., Lv, X., Irwin, D.M., Savolainen, P., Wu, C.-I., Zhang, Y., 2013. The genomics of selection in dogs and the parallel evolution between dogs and humans. Nat Commun 4, 1860. https://doi.org/10.1038/ncomms2814

Wang, G.-D., Zhai, W., Yang, H.-C., Wang, L., Zhong, L., Liu, Y.-H., Fan, R.-X., Yin, T.-T., Zhu, C.-L., Poyarkov, A.D., Irwin, D.M., Hytönen, M.K., Lohi, H., Wu, C.-I., Savolainen, P., Zhang, Y.-P., 2016. Out of southern East Asia: the natural history of domestic dogs across the world. Cell Res 26, 21–33. https://doi.org/10.1038/cr.2015.147

Wayne, R. K., Vilà, C. 2003. Molecular Genetic Studies of Wolves. In Wolves: behavior, ecology, and conservation. L. D. Mech and L. Boitani, ed. (University of Chicago Press), pp. 218–238.

Werhahn, G., Senn, H., Ghazali, M., Karmacharya, D., Sherchan, A.M., Joshi, J., Kusi, N., López-Bao, J.V., Rosen, T., Kachel, S., Sillero-Zubiri, C., Macdonald, D.W., 2018. The unique genetic adaptation of the Himalayan wolf to high-altitudes and consequences for conservation. Global Ecology and Conservation 16, e00455. https://doi.org/10.1016/j.gecco.2018.e00455

Wozencraft, W. C. 2005. Order Carnivora. In Mammal species of the world, D. E. Wilson and D. M. Reeder, ed. (Smithsonian Institution Press), pp. 532–628.

Zhang, C., Rabiee, M., Sayyari, E., Mirarab, S., 2018. ASTRAL-III: polynomial time species tree reconstruction from partially resolved gene trees. BMC Bioinformatics 19, 153. https://doi.org/10.1186/s12859-018-2129-y

Zhang, W., Fan, Z., Han, E., Hou, R., Zhang, L., Galaverni, M., Huang, J., Liu, H., Silva, P., Li, P., Pollinger, J.P., Du, L., Zhang, X., Yue, B., Wayne, R.K., Zhang, Z., 2014. Hypoxia Adaptations in the Grey Wolf (Canis lupus chanco) from Qinghai-Tibet Plateau. PLOS Genetics 10, e1004466. https://doi.org/10.1371/journal.pgen.1004466

