## Supplementary material for "Conservation implications of elucidating the Korean wolf taxonomic ambiguity through whole-genome sequencing": Suplemental information

Table S1. Resequenced wolf samples.

| Project ID | Museum ID | Date | Museum of origin | Species | Country | Sex | Tissue | Average whole genome coverage |
|---|---|---|---|---|---|---|---|---|
| Pyongyang Zoo wolf | IN149 | 2005 | Seoul Grand Park, South Korea | *Canis lupus* | South Korea | Female | Blood | 25.2989 |
| Historical Korean wolf | / | Before 1945 | Kyungpook University Museum, Seoul, South Korea | *Canis lupus* | South Korea | Female | skin | 7.2511 |
| MW486 | V25-RU | 2000 | Mikael Åkesson & Jouni Aspi (SLU) | *Canis lupus* | Russia | Male | skin | 5.811174736 |
| MW524 | 3 | 2015 | Morten Skage (Oslo, Norway) | *Canis lupus* | Russia | Female | Extracts | 5.784092198 |
| MW536 | 15 | 2015 | Morten Skage (Oslo, Norway) | *Canis lupus* | Russia | Female | Extracts | 5.732749185 |
| MW538 | 17 | 2015 | Morten Skage (Oslo, Norway) | *Canis lupus* | Russia | Male | Extracts | 5.850073506 |
| MW574 | 24 | 2019 | Boldgiv Bazartseren | *Canis lupus* | Mongolia | Female | Cheek tissue | 5.770702451 |
| MW588 | 43 | 2019 | Boldgiv Bazartseren | *Canis lupus* | Mongolia | Female | Fresh skin | 5.495638571 |
| MW561 | 4 | 2018 | Boldgiv Bazartseren | *Canis lupus* | Mongolia | Female | Muscle tissue | 5.481997786 |
| 750021A | 750021 | modern | Yekaterinburg Museum | *Canis lupus* | Russia | Female | Muscle tissue | 5.460293376 |
| 750115A | 750115 | modern | Yekaterinburg Museum | *Canis lupus* | Kazakhstan | Male | Muscle tissue | 6.431657396 |
| MNG-A2 | A70094 | Birth 25/3/2007 | Zoo Zürich | *Canis lupus* | Germany | Male | Muscle tissue | 7.64 |

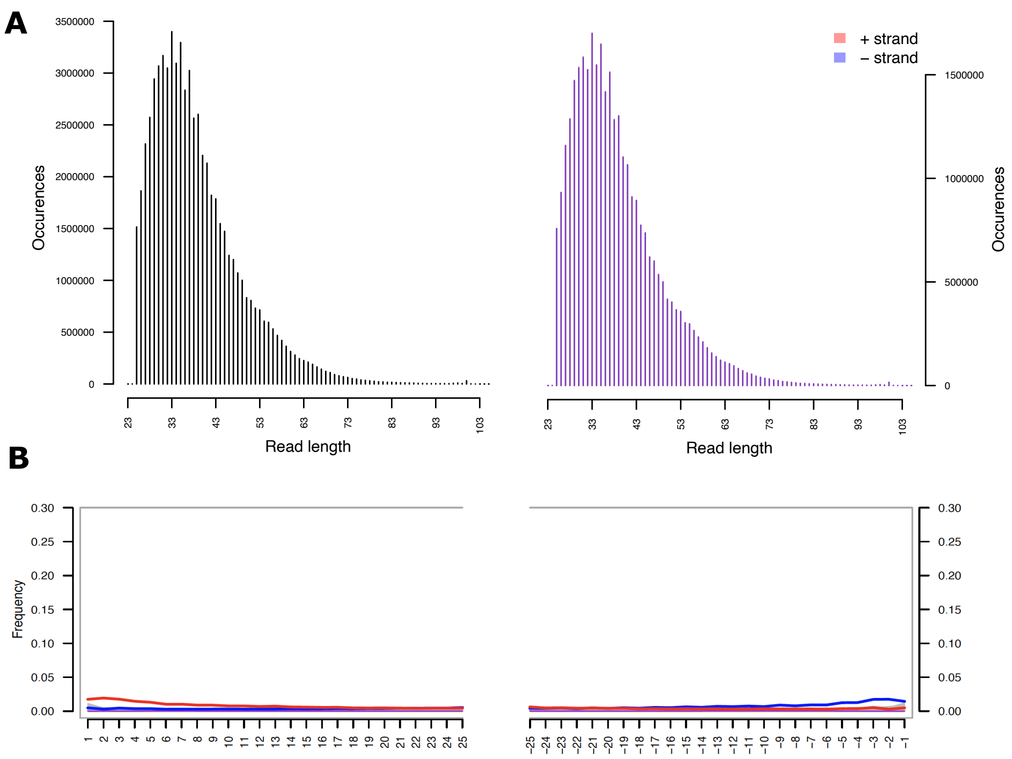


**Figure S1. DNA damage assessment of historical Korean wolf data.**
A) Sequence reads length distribution. Predominantly, short reads were obtained for the sequencing of the stuffed historical specimen. B) DNA damage pattern shown by the increase of C to T and G to A substitutions at the read ends. A low percentage of DNA damage can be observed in the historical Korean wolf sequenced reads.

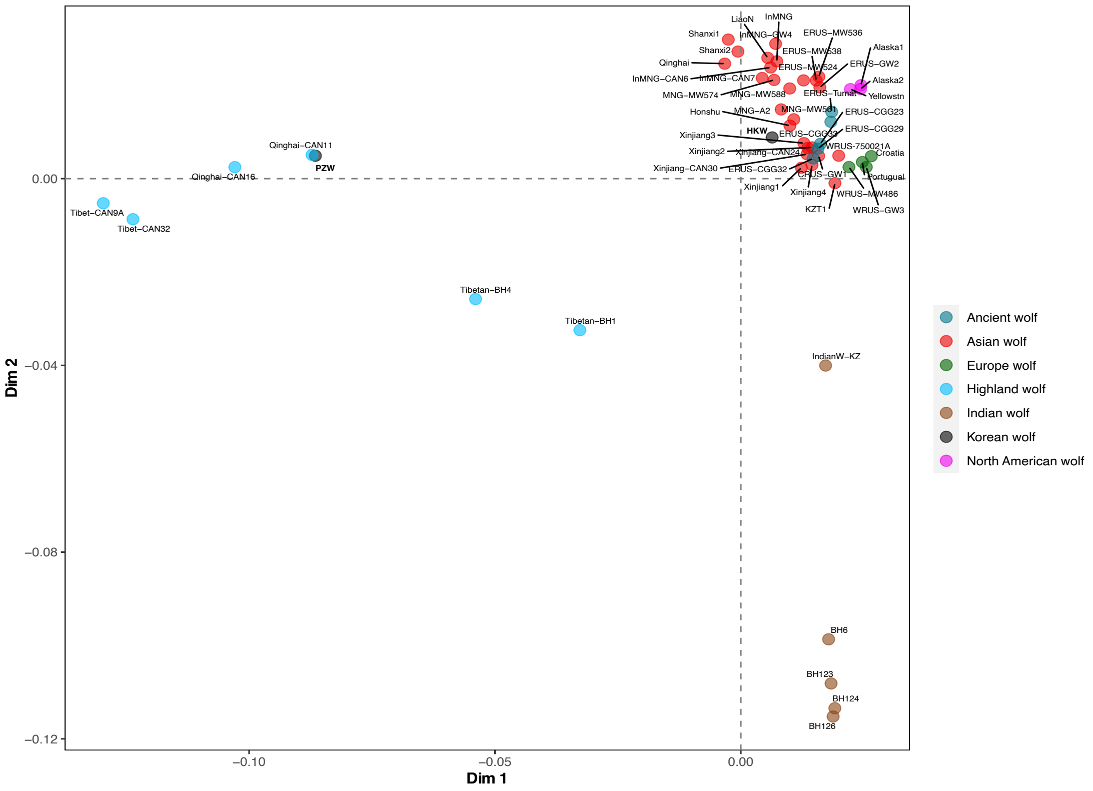


**Figure S2. Extended MDS plot result. Related to Figure 1A.**

MDS plot including just wolf genomes to explore in more detail the population structure among wolf. The analysis was performed using a pseudo-haploid dataset of 3,284,758 transversion sites.

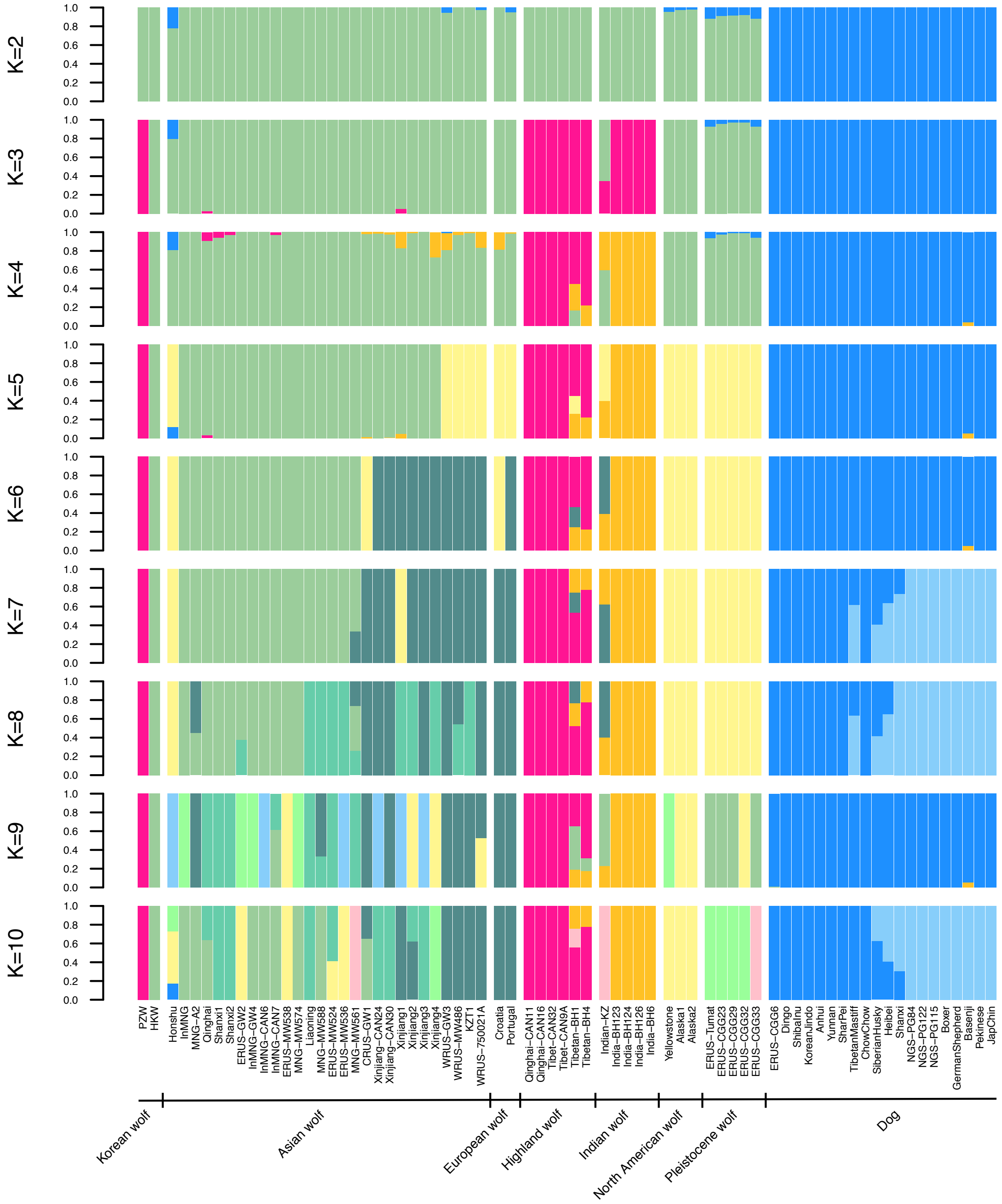


**Figure S3. Extended admixture graphs estimated for 2 to 10 ancestry components (K). Related to Figure 1C.**

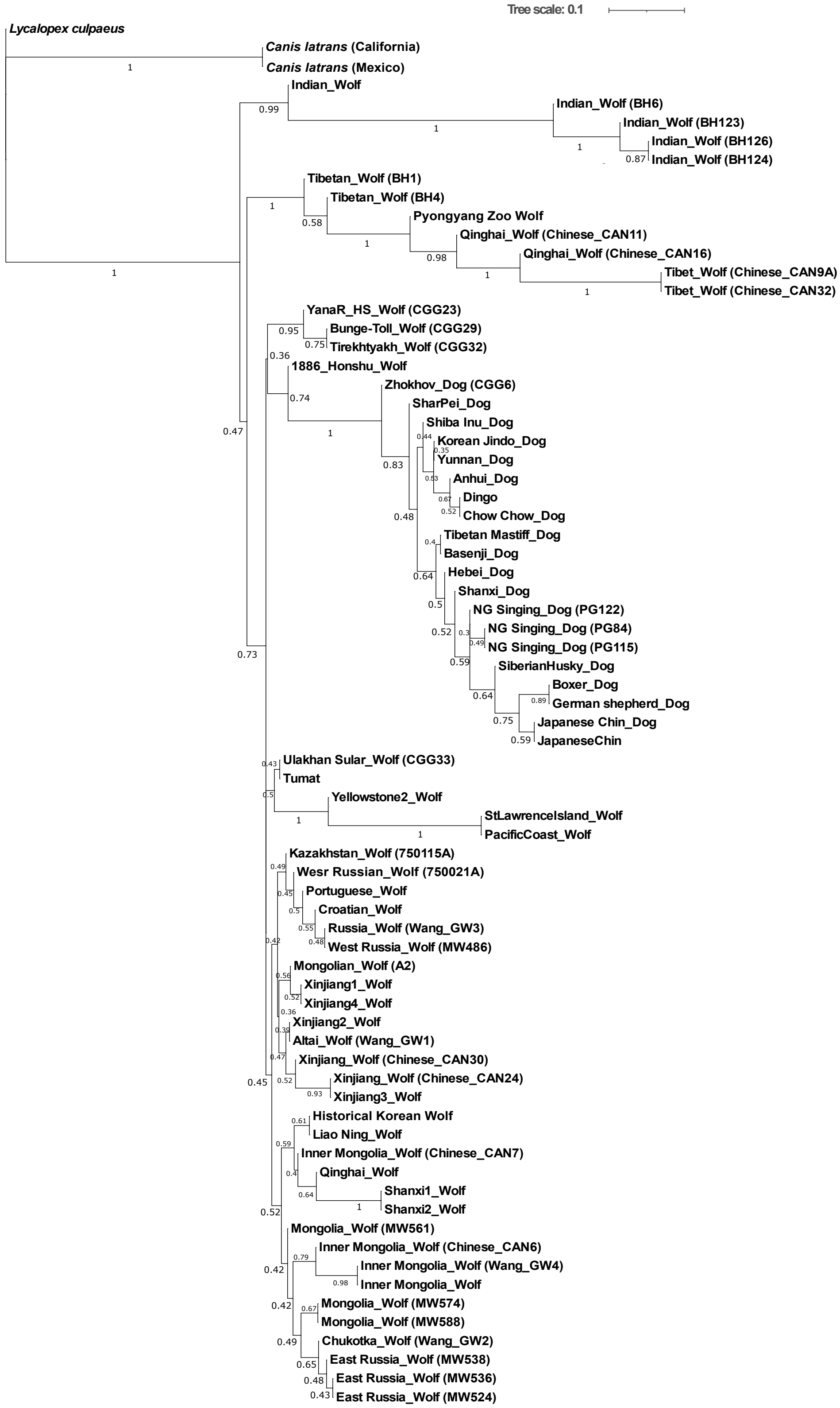


**Figure S4. Phylogenetic relationships estimated by genomic data.**
Maximum likelihood phylogeny built on 1000 concatenated trees. Bootstrap values are shown for each internal branch.

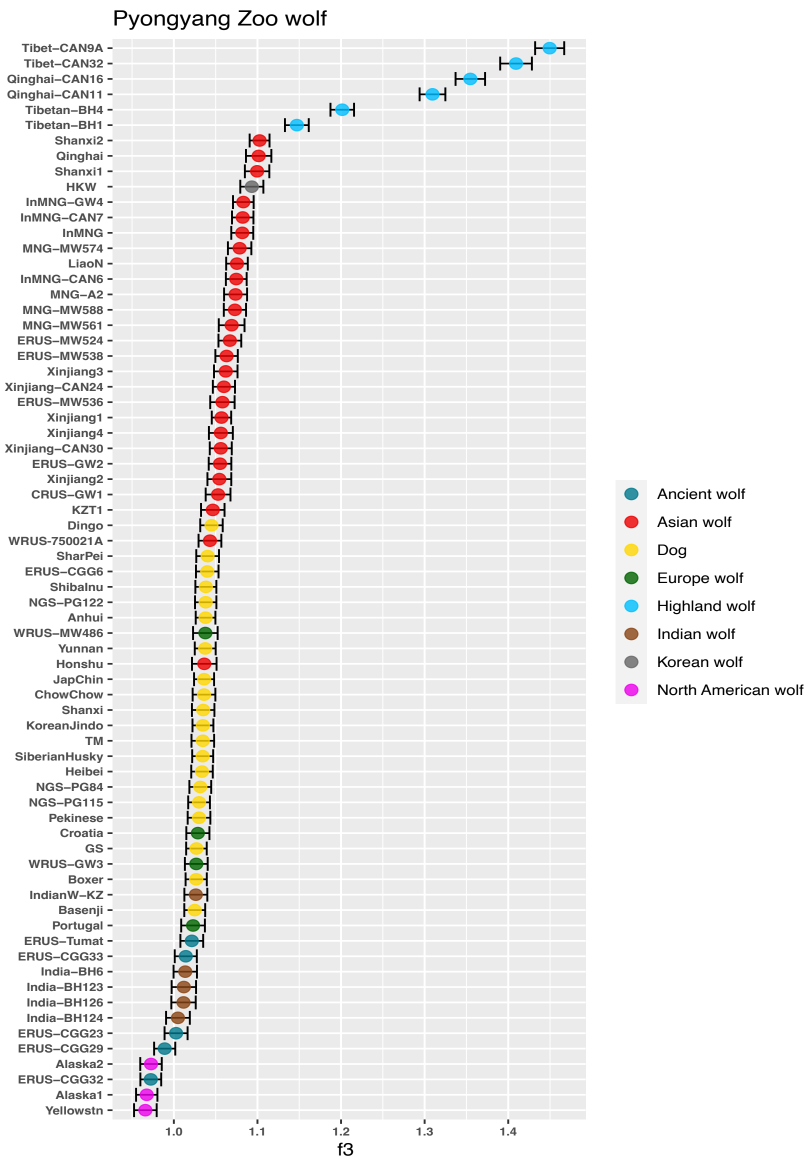


**Figure S5. Assessment of shared genetic drift between the Pyongyang Zoo wolf and other wolves and dogs in the dataset.**

To test the genetic affinities of the Pyongyang Zoo wolf we performed an outgroup *f3-statistics* analysis using the coyotes as outgroup.

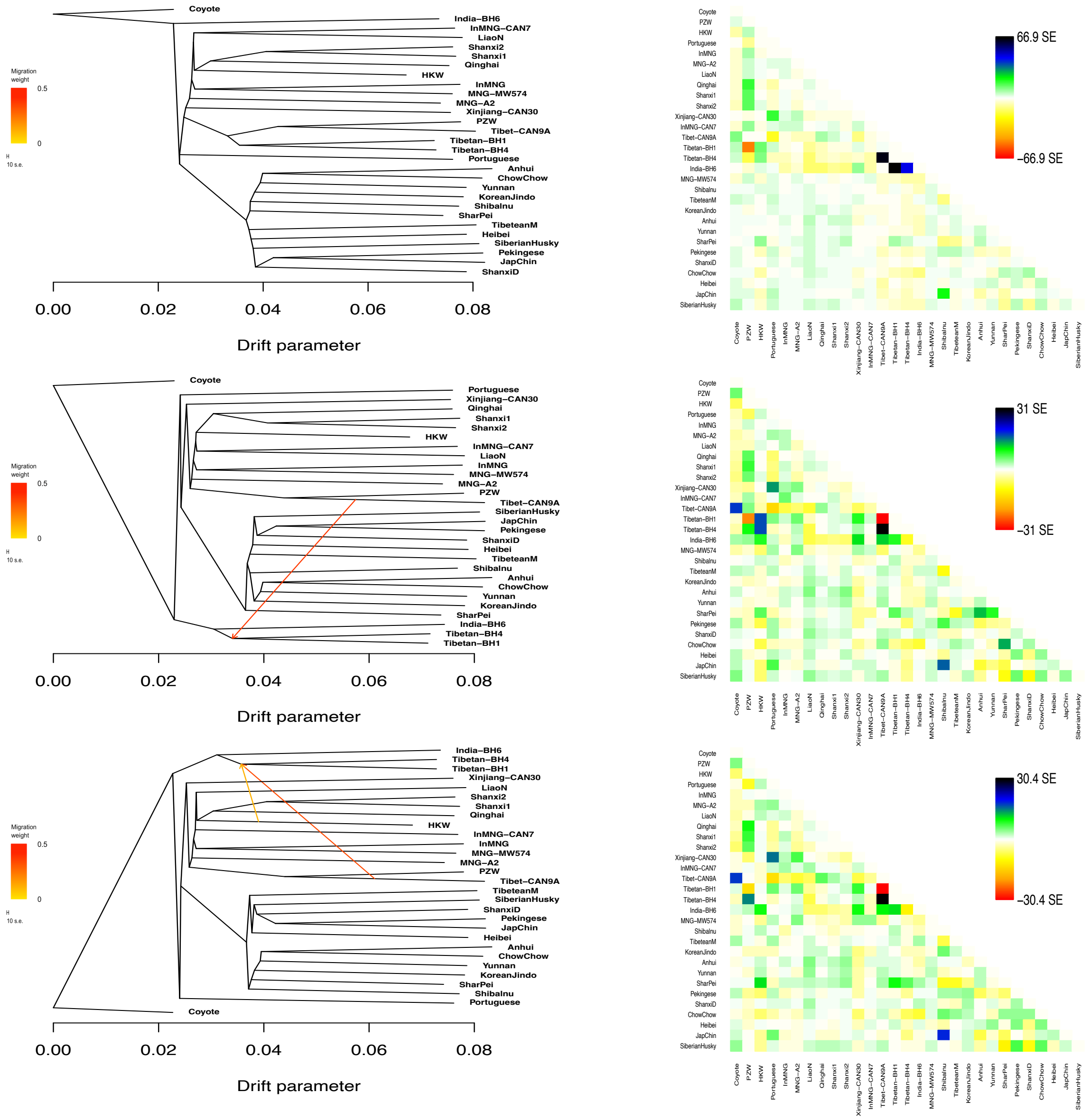


**Figure S6. Extended Treemix admixture graphs and residual plots. Related to Figure 2C.** Treemix graphs for 0 to 2 migration edges and residual plots from the fit of the model to the data. The analyses were run using an SNP panel of 1,904,538 transversion sites, excluding sites with missing data.

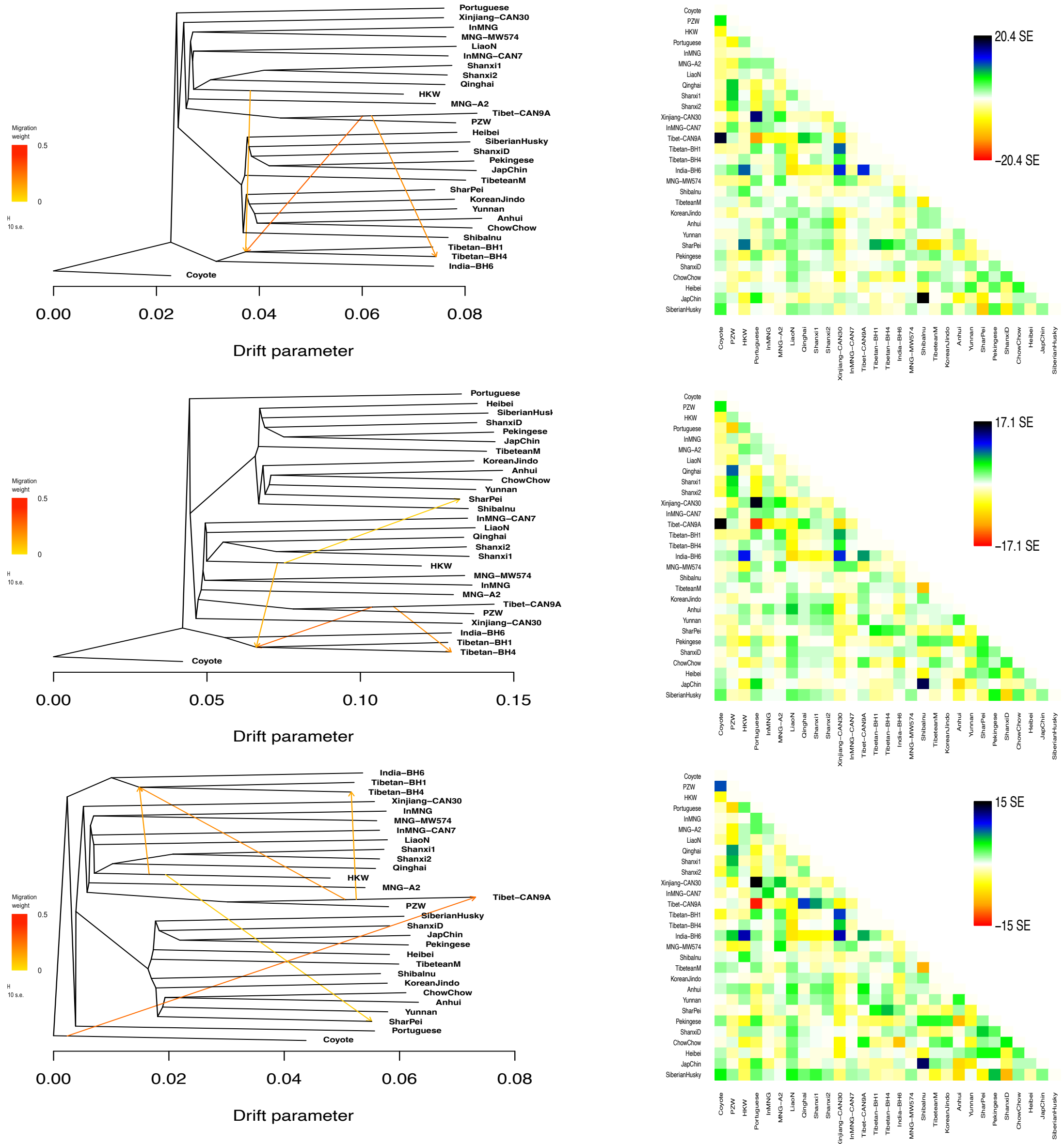


**Figure S7. Extended Treemix admixture graphs and residual plots. Related to Figure 2C.** Treemix graphs for 3 to 5 migration edges and residual plot from the fit of the model to the data.

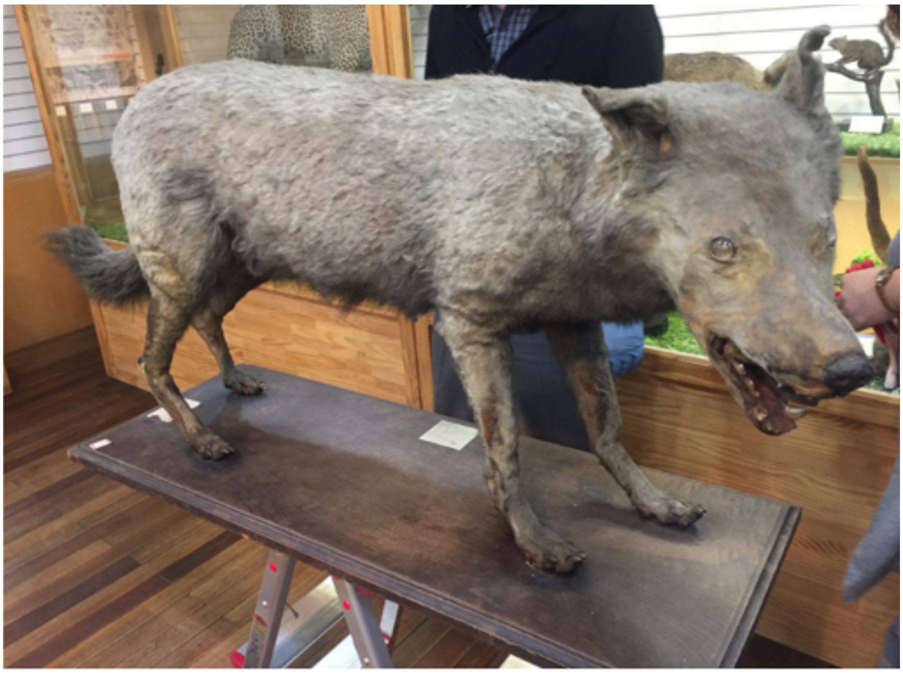

**Figure S8. Stuffed historical Korean wolf specimen sequenced in this study.**
